# FastGxC: Fast and Powerful Context-Specific eQTL Mapping in Bulk and Single-Cell Data

**DOI:** 10.1101/2021.06.17.448889

**Authors:** Lena Krockenberger, Andrew Lu, Mike Thompson, Cuining Liu, M Grace Gordon, Amanda Ramste, Ivan Carcamo-Orive, Joshua W. Knowles, Andy Dahl, Chun Jimmie Ye, Noah Zaitlen, Brunilda Balliu

## Abstract

Context-specific eQTLs mediate genetic risk for complex diseases. However, limitations in current methods for identifying these eQTLs have hindered their comprehensive characterization and downstream interpretation of disease-associated variants. Here, we introduce FastGxC, a method to efficiently and powerfully map context-specific eQTLs by leveraging the correlation structure in genomic studies with repeated sampling, e.g., single-cell RNA-seq studies. Using simulations, we demonstrate that FastGxC is up to nine times more powerful and 10^6^ times faster than existing approaches, reducing computation time from years to minutes. We applied FastGxC to bulk multi-tissue (N=698) and single-cell PBMC (N=1,218) RNA-seq datasets, generating comprehensive tissue- and cell-type-specific eQTL maps. These eQTLs exhibited up to four-fold enrichment in open chromatin regions from matched contexts and were twice as enriched as standard context-specific eQTLs, highlighting their biological relevance. Furthermore, we examined the relationship between context-specific eQTLs and complex human traits and diseases. FastGxC improved precision in identifying relevant contexts for each trait by three-fold and expanded candidate causal genes by 25% in cell types and 6% in tissues compared to standard eQTLs. In summary, FastGxC provides a powerful framework for mapping context-specific eQTLs, advancing our understanding of gene regulatory mechanisms underlying complex human traits and diseases.

## 1 Introduction

Over the past 15 years, genome-wide association studies (GWAS) have identified tens of thousands of genetic variants linked to complex traits and diseases [1]. A majority of these variants reside in non-coding regions, often overlapping DNA regulatory elements [2], which suggests their functional effects are mediated through transcriptional regulation [2–5]. This observation has driven significant efforts to identify expression quantitative trait loci (eQTLs) — genetic variants associated with gene expression changes — and use them to link GWAS variants to their regulatory targets [6–15]. Despite these efforts, only 21% of GWAS variants, on average per trait, overlap with known cis eQTLs from bulk tissues [12], underscoring a persistent gap between genetic associations and regulatory function [16–18].

A key factor for this missing regulation is the context-specific nature of many disease-relevant eQTLs [16, 18], which often appear only in specific tissues [12], cell types [10, 14, 15], or environmental conditions [19–23], making them difficult to detect. In contrast, broadly shared eQTLs, while easier to detect, are less enriched for GWAS variants [12, 14], likely due to negative selection [16, 18]. Another major factor is that many bulk [8, 12] and all single cell RNA-Sequencing (RNA-Seq) studies rely on repeated sampling, where the same donor provides samples across multiple contexts. While this design minimizes experimental variability, it introduces intra-individual correlation, which, if unaccounted for, inflates type I error rate to identify an eQTL and reduces the power to test if the eQTL is context-specific.

Several methods have been developed to identify context-specific eQTLs in studies with repeated sampling (see Table S1). These methods fall into two broad categories. The first comprises approaches that jointly analyze data across contexts and test for context-specific eQTLs by incorporating a genotype-by-context (GxC) interaction term. Note that, while we refer to eQTLs with significant GxC effects as context-specific, in alignment with common genomics terminology [24, 25], the more precise term would be context-dependent eQTLs. This includes (generalized) linear mixed model (LMM)-based methods [24–29], which model the GxC effect linearly, and methods that capture non-linear GxC effects [30]. To account for repeated measurements, these methods include a random effect for the individual or cell. While powerful, their mixed model framework makes them computationally intensive for large eQTL studies. This challenge is particularly exacerbated when modeling all cell types jointly. Additionally, some of these methods [24, 30] infer latent cellular contexts, further increasing computational costs.

The other category includes methods that follow a two-step process: first, they map eQTLs separately in each context (context-by-context; CxC), then they define context-specificity by post hoc examination of eQTL summary statistics across contexts [10, 12, 15, 31–33]. While CxC approaches are fast, particularly those developed for (pseudo)-bulk data [10, 12, 15, 33], they have major limitations. First, CxC approaches can be significantly underpowered because they do not fully leverage all available data. Second, many rely on ad hoc definitions of context-specificity based on subjective thresholds of effect size differences between contexts [33, 34] or the significance of an eQTL in a single context [10, 12, 15]. These definitions can lead to both false-positive context-specificity (e.g., when effects in certain contexts fail to reach significance due to chance or uneven power across contexts) and false-negative context-specificity (e.g., when an eQTL is shared across contexts but still shows GxC interaction effects). Taken together, these limitations constrain the interpretation of disease-associated variants, as current methods fail to fully capture the complexity of context-specific regulatory variation.

To address these challenges, we introduce FastGxC, a novel method that efficiently maps context-specific eQTLs while accounting for repeated sampling. In brief, FastGxC decomposes gene expression into context-shared and context-specific components and estimates genetic effects on these components using linear regression. We show analytically and empirically that FastGxC’s eQTL effect estimates can be viewed as computationally efficient reparametrizations of those obtained through CxC and LMM-GxC approaches. FastGxC has several key advantages over previous methods. First, it directly maps specific eQTLs without the need for post hoc analyses or arbitrary thresholds. Second, by accounting for intra-individual correlation, it adjusts for background noise and confounding factors unrelated to the context of interest, e.g., sex, age, population stratification, or sequencing batch [35–37], maximizing power to detect context-specific eQTLs (Figure S1). Third, FastGxC leverages ultra-fast implementations of linear regression models, similar to those used in CxC eQTL mapping approaches [38–40], which reduce computational time from years to minutes. FastGxC can work on any continuous molecular phenotype and its output integrates naturally with methods developed to improve the statistical power of eQTL mapping, such as mash [34].

We first show in simulations that FastGxC is as powerful as the LMM-GxC approachs but orders of magnitude faster. Both approaches significantly outperform CxC-based methods to map context-specific eQTLs. We then applied FastGxC to multi-tissue bulk RNA-Seq data from the GTEx Consortium[12] (N=698 individuals) and single-cell peripheral blood mononuclear (PBMC) RNA-Seq data from the CLUES [14] (N = 237) and OneK1K[15] (N = 981) cohorts, which we meta-analyze, to produce comprehensive tissue- and cell type-specific eQTL maps across 49 tissues and 8 PBMC cell types. FastGxC context-specific eQTLs show up to four-fold enrichment in open chromatin regions from matched contexts and are twice as enriched as standard context-specific eQTLs, highlighting their biological relevance. We further examine their relationship with complex traits and diseases, showing that FastGxC eQTLs improve precision in identifying relevant GWAS contexts by three-fold and expand candidate causal genes by 25% in cell types and 6% in tissues compared to standard eQTLs. In addition, FastGxC context-specific eQTLs show a 1.2-fold increase in colocalization with complex traits compared to shared eQTLs, providing evidence that context-specific regulation helps explain regulatory mechanisms of complex diseases that remain unaccounted for by shared eQTLs. In summary, FastGxC provides a powerful framework for constructing context-specific eQTL maps, offering key insights into the gene regulatory mechanisms underlying complex human diseases.

## 2 Results

### FastGxC method overview

We illustrate the FastGxC method using tissues as contexts (Figure 1A), but the method can be applied to any set of discrete contexts, for example, cell types [10, 14, 15] or environmental stimuli, sampled across overlapping individuals. FastGxC works in three steps. First, for each individual *i* (*i* = 1, …, *N*) and context *c* (*c* = 1, …, *C*), FastGxC decomposes the expression of each gene (*E*_*ic*_) into two components: a shared component 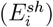, representing the average expression across contexts, and a context-specific component 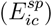, representing the residual expression in the contexts after subtracting the shared component (Figure 1A - Decomposition step), i.e., 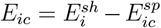. This decomposition, analogous to repeated-measures ANOVA, removes shared eQTLs effects and shared noise from the context-specific components, thereby increasing power to detect context-specific eQTLs (Figure S1) [41].

**Figure 1.**
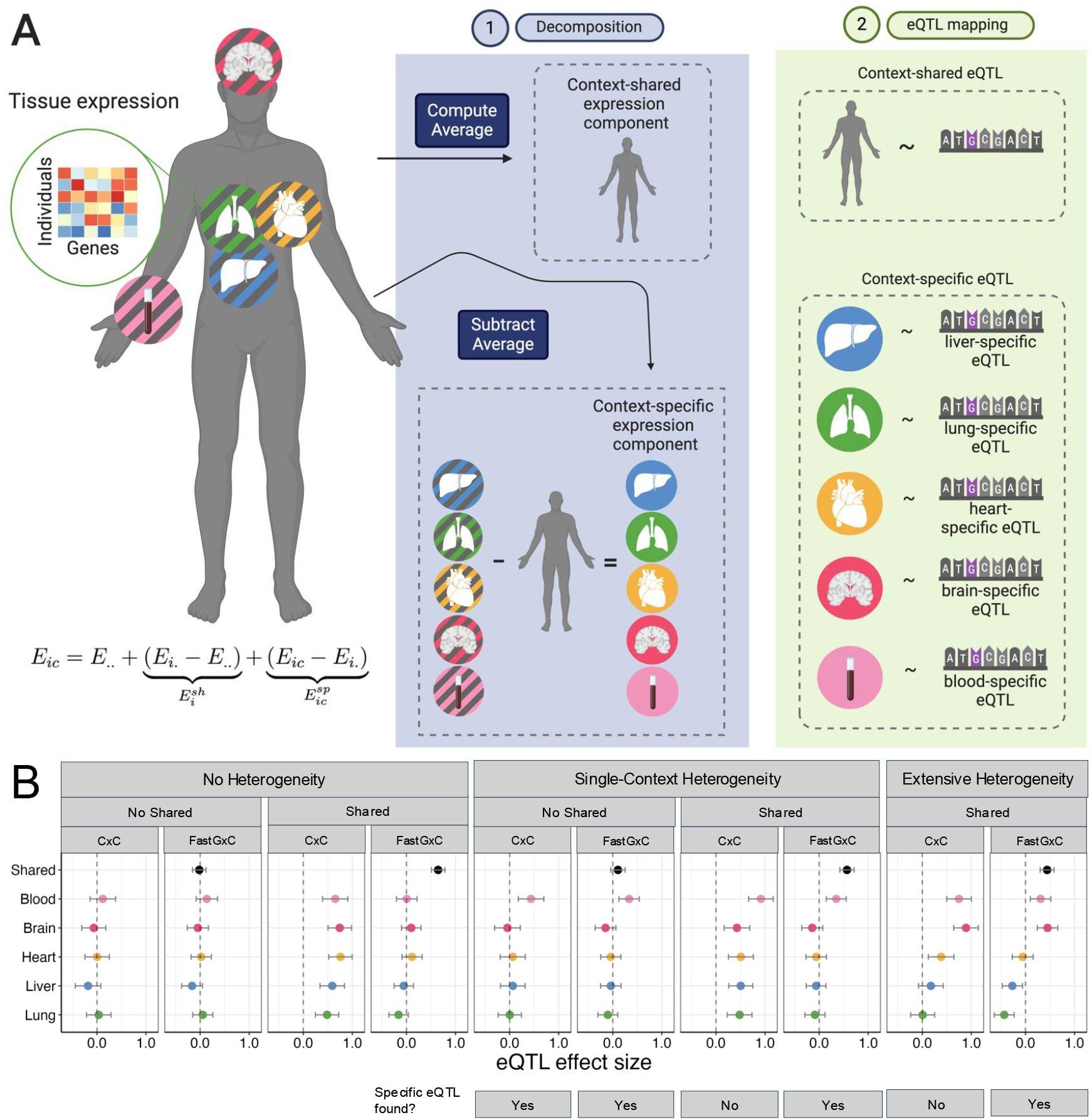
Overview of the FastGxC method and toy examples of eQTLs. **A.**FastGxC decomposes gene expression for each individual into a context-shared component and context-specific components (Step 1). It then estimates both the shared eQTL effect across contexts and the context-specific eQTL effects within each context by regressing genotypes on these components (Step 2). **B**. Toy examples of eQTLs. Y axis and color represent the context and x axis lists the eQTL effect. The first example represents a scenario with no eQTLs in any tissue and, thus, no shared or specific eQTLs. The second example represents a scenario with equal eQTL effects across all tissues, corresponding to a scenario with a shared eQTL but no specific eQTLs. The third, fourth, and fifth examples corresponds to a scenario with an eQTL in which a single context (e.g., blood) or multiple contexts drive the effect size heterogeneity.

Next, for each gene–cis-SNP pair, FastGxC estimates a shared eQTL effect (*β*^*sh*^) and *C* specific eQTL effects 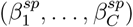 by regressing the genotype at the cis-SNP on the shared expression component and each of the *C* context-specific components (Figure 1A - eQTL mapping step). This step employs ultra-fast implementations of linear regression models optimized for eQTL mapping [38, 39], enabling computational efficiency comparable to standard eQTL mapping methods. The shared and context-specific effects represent a reparametrization of the eQTL effects obtained from conventional context-by-context eQTL mapping (*β*_1_,…, *β*_*C*_)(Figure 1B and S16). Specifically, the shared effect corresponds to the mean eQTL effect size across contexts, i.e., 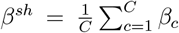 while the context-specific effects capture the residual eQTL effects in each context after accounting for the shared effect, i.e., 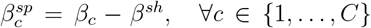. This decomposition separates the pleiotropic (shared) effect of an eQTL across all contexts from the context-specific effects, enabling clearer interpretation of context-specific genetic effects. Because CxC itself can be viewed as a reparametrization of the LMM-GxC framework, FastGxC provides a computationally efficient reparametrization of LMM-GxC. Full details of the analytical derivation are provided in the Online Methods and Supplementary Text.

Finally, to account for multiple testing across genes, SNPs, and contexts, FastGxC employs the hierarchical False Discovery Rate (FDR)-controlling procedure implemented in [42] (Online Methods and Figure S2). FastGxC defines a gene-SNP pair as an eQTL if the SNP has a significant effect on the shared or any of the specific components of gene expression, i.e., if the global null hypothesis 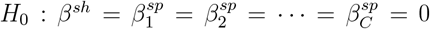 is rejected. If an eQTL is detected, FastGxC defines a context-specific eQTL as a SNP with a significant effect on at least one of the specific components of expression of the gene, i.e., if the global null hypothesis 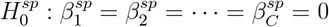 is rejected.

This global test directly identifies context-specific eQTLs, eliminating the need for post hoc analyses or arbitrary thresholds. Finally, if significant eQTL effect size heterogeneity is detected, FastGxC conducts *C* marginal tests to determine the specific context(s) driving the observed heterogeneity, i.e., 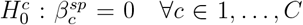. Note that these tests do not specifically flag the contexts with non-zero eQTL effects; rather, they detect contexts whose effect sizes deviate significantly from the shared effect. To illustrate how FastGxC identifies different patterns of context-specificity, Figure 1B shows toy examples of eQTLs with varying patterns of effects across contexts.

The first two panels (“No Heterogeneity”) depict scenarios under the null hypothesis of no eQTL effect size heterogeneity across contexts, with either a shared eQTL effect (“Shared”) or no shared eQTL effect (“No Shared”). FastGxC does not classify either of these eQTLs as context-specific, as there is no significant heterogeneity of their effects across contexts, but it would classify the second scenario as a (shared) eQTL. The remaining panels illustrate scenarios under the alternative hypothesis of eQTL effect size heterogeneity. These include heterogeneity driven by a single context (“Single-context Heterogeneity”) or heterogeneity spanning all contexts (“Extensive Heterogeneity”). FastGxC identifies all these scenarios as context-specific eQTLs, irrespective of the presence of a shared eQTL effect. By contrast, the commonly used CxC approach defines context-specific eQTLs as variants with significant eQTL effects in only a single context. As a result, this approach would classify only the first alternative scenario (“Single-context Heterogeneity - No Shared”) as a context-specific eQTL and would overlook more complex patterns of heterogeneity, such as cases where heterogeneity exists alongside a shared effect or where heterogeneity is distributed across multiple contexts, highlighting its limitations compared to FastGxC.

### FastGxC outperforms existing methods in simulation studies

We used a series of simulated scenarios to evaluate the performance of FastGxC to detect an eQTL and determine if the eQTL effect is context-specific as a function of intra-individual residual correlation (see Online Methods and Table S2). In each scenario, we varied the number of individuals and contexts and the proportion of missing expression data to reflect those in GTEx [12] and the OneK1K cohort [15], two of the largest bulk and single cell RNA-Seq studies. The performance of FastGxC was systematically compared to three commonly used approaches: (1) the CxC approach, which performs context-by-context eQTL mapping and defines a context-specific eQTL as a variant with a significant effect in a single context, (2) MetaTissue [43], a multi-tissue eQTL mapping method that combines mixed models and meta-analysis, and defines a context-specific eQTL as a variant with a posterior probability greater than 0.9 of having an effect present in exactly one context, and (3) the linear mixed model (LMM-GxC) approach, which includes a random intercept for individuals to account for intra-individual residual correlation and defines a context-specific eQTL based on the significance of the genotype-by-context (GxC) interaction term (see Online Methods). To illustrate the impact of ignoring intra-individual correlation on the identification of context-specific eQTLs, we also include performance of a linear model with a GxC interaction term (LM-GxC) but no random intercept. Note that due to the large computational burden of MetaTissue, we did not obtain results for all scenarios with larger sample size (N=698).

We first evaluated the global type I error rates of each method for detecting an eQTL (Figure 2A and S3) and for testing whether an eQTL is context-specific (Figure 2A and S4). FastGxC is well-calibrated across all tested scenarios and for both tests. LMM-GxC is generally calibrated but becomes inflated in settings with low sample size and high missing data rates. As expected, both the CxC and LM-GxC approaches, which do not account for intra-individual correlation, are miscalibrated (Figure S3). As intra-individual correlation increases, LM-GxC becomes increasingly inflated for eQTL detection and increasingly conservative for testing context-specificity. The CxC approach, by contrast, remains mostly calibrated when testing for the presence of an eQTL. However, depending on sample size, missing data rate, and the presence or absence of a shared eQTL, CxC becomes either increasingly conservative (Figure 2A) or anti-conservative (Figure S4) when testing for (single-)context-specificity. Finally, MetaTissue is consistently conservative in scenarios with low sample size (Figure S3) when testing for the presence of an eQTL. However, when evaluating context specificity, MetaTissue becomes anticonservative or conservative, depending on whether a shared eQTL effect is present (Figure S4).

**Figure 2.**
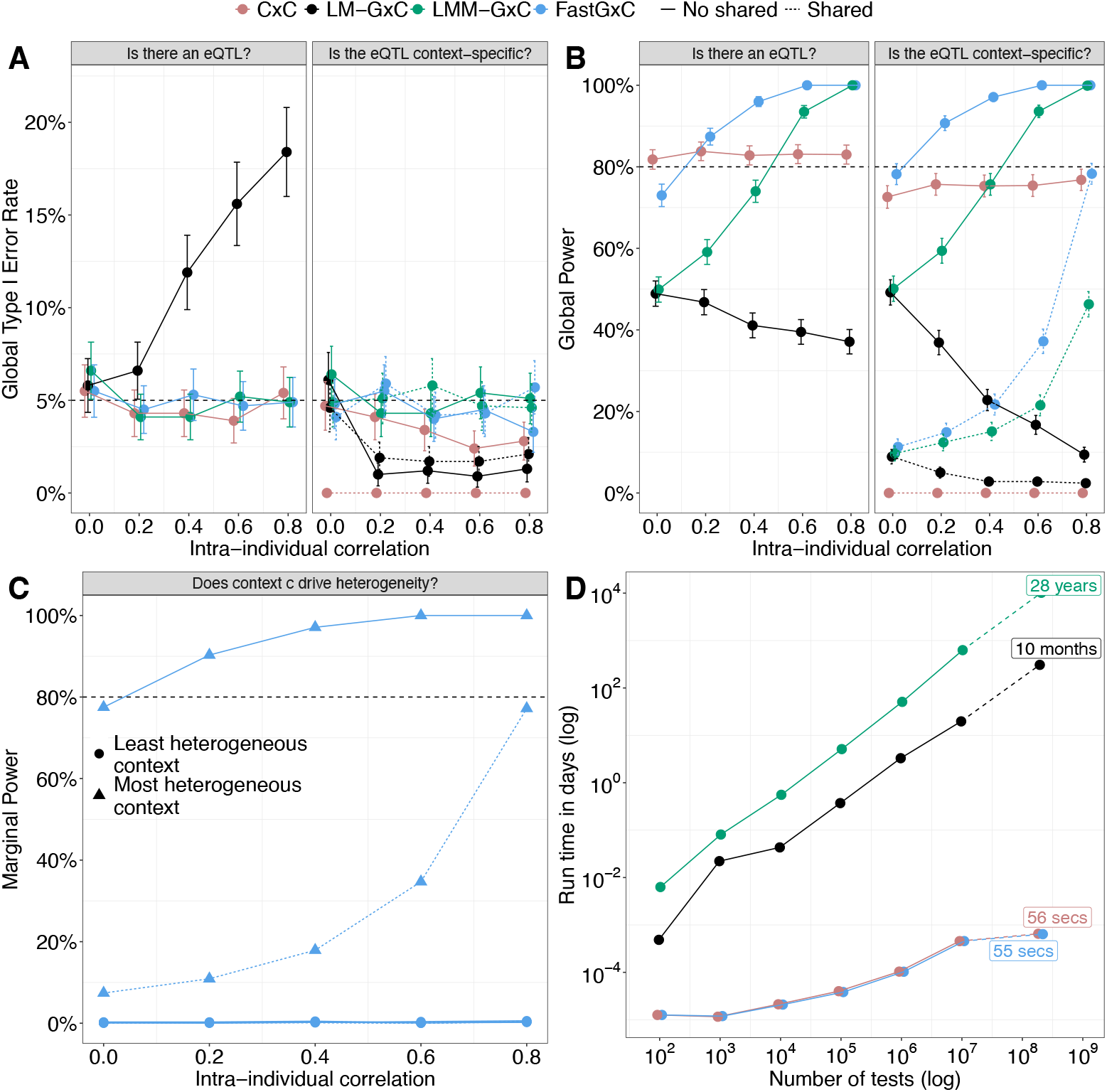
FastGxC outperforms existing methods in simulated data. **A-B.** Global Type I error rate (A) and global and marginal power (B and C) for detecting an eQTL (A and B - left panel), testing for context-specificity of its effect (A and B - right panel), and identify which context drives the heterogeneity (C) under the no heterogeneity (A) and single-context heterogeneity (B, C) scenarios (Figure 1B) across different levels of intra-individual correlation (rows). For effect sizes in each scenario, see Table S2. For power under the two-context and extensive heterogeneity scenarios, see Figures - S10. For marginal Type I error rates and marginal power under the two-context and extensive heterogeneity scenarios see Figures - S14. Figure panels A-C show results from simulations with 698 individuals and 49 contexts and GTEx missing data patterns (63%). For results without or less missing data, lower sample size, and fewer contexts see Figures - S14. **D**. Run time for all methods for varying number of tests performed in a sample size of 250 individuals (average sample size across tissues in GTEx). See Figure for sample size of 1,000 individuals. Last points reflect projected run time for entire GTEx data-set - 50 contexts, 25K x 3M tests, and 250 samples per context. Analyses were run on 8 cores on a 2.70 GHz Intel Xeon Gold Processor on the UCLA Hoffman2 Computing Cluster.

Next, we evaluated the global power of each method to identify an eQTL. Among the calibrated methods, FastGxC and LMM-GxC exhibit complementary strengths. FastGxC is generally more powerful when eQTL effect size heterogeneity is strong or driven by a few contexts (Figure 2B, S6–S8), as it leverages the Simes’ method to combine p-values. LMM-GxC is more powerful when heterogeneity is weak but spread across many or all contexts (Figure S9–S10), due to its reliance on the likelihood ratio test (LRT). The CxC and MetaTissue approaches are less powerful than FastGxC to identify an eQTL for all scenarios with non-zero intra-individual correlation, with the power advantage of FastGxC increasing as correlation rises. The LM-GxC method is miscalibrated for eQTL detection; therefore, we do not report or discuss its power to map an eQTL.

For all methods, power to detect an eQTL also depends on whether variability in expression is explained by shared or context-specific effects. When the shared eQTL effect explains all (“No heterogeneity”, Figure S5) or most of the expression variability (“Single-context Heterogeneity - Shared”, Figure S6), power to identify an eQTL declines as intra-individual correlation increases, whereas the opposite occurs when the specific eQTL effect explains all or most variability (“Single-context Heterogeneity - No shared”, Figure S6). In scenarios with intermediate shared and specific effects, power to identify an eQTL follows a U-shaped relationship with intra-individual correlation (“Extensive heterogeneity”, Figure S9).

We next examined power to assess eQTL context specificity. For both FastGxC and LMM-GxC, power increases as intra-individual correlation increases, regardless of whether a shared eQTL effect is present. Notably, FastGxC is more powerful than the CxC and MetaTissue approaches even in the single-context heterogeneity scenario without a shared effect—the scenario in which these approach are specifically used. As expected from its performance under the null, the LM-GxC method loses power to test for eQTL context specificity as intra-individual correlation increases (Figures 2B, S6–S10).

Then, to evaluate the ability of FastGxC to identify specific contexts driving effect size heterogeneity, i.e. contexts most different from the shared effect, we examined the marginal type I error rate and power per context. Under the null hypothesis (“No heterogeneity”), FastGxC is calibrated for each context (FDR ≤5%, Figure S11). Under the alternative hypothesis, power was highest for contexts with effect sizes farthest from the shared effect and increased with intra-individual correlation (Figures 2C, S12–S14). For example, in single-context heterogeneity scenarios, FastGxC accurately identifies the context with the non-zero (“No shared” scenario) or single strongest eQTL effect (“Shared” scenario) (Figures 2C and S12).

In addition, we assessed FastGxC’s parameter estimation accuracy by evaluating its ability to estimate the shared (i.e., *β*^*sh*^) and specific eQTL effect sizes (i.e., 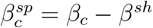 as well as the overall eQTL effect sizes in each context (i.e., 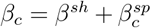; Figure S16-S15). FastGxC provided unbiased estimates for the shared, specific, and overall eQTL effect sizes under conditions with no missing data or with missing data levels typical of single-cell RNA-Seq studies such as OneK1K and CLUES (approximately 5%). When the proportion of missing data was high (mean of 63% and up to 84% in some contexts), FastGxC estimates remained largely unbiased, with only slight deviations from the true effect in contexts with high missing rate.

Finally, we benchmarked the computational costs of FastGxC against other approaches. To obtain practical run-times, we used study parameters from GTEx, i.e., approximately 50 contexts and an average of 250 individuals per context, while varying the number of tests performed (Figure 2D). When extrapolated to the entire GTEx dataset, which involves 200 million tests for 25,000 genes and 3 million SNPs, we estimated that LMM-GxC and LM-GxC would require approximately 30 years and 10 months, respectively, to complete. In contrast, CxC and FastGxC completed the same task in under one minute on average (based on 100 iterations). Even at a larger scale with 1,000 individuals, FastGxC remained computationally efficient, completing all tests in approximately five minutes, whereas LMM-GxC was estimated to take over 500 years (Figure S17). MetaTissue was not included in the runtime analysis due to its substantial computational burden, which exceeds that of LMM-GxC.

### Context-specificity of eQTLs is widespread across tissues and PBMC cell types

To evaluate performance in bulk tissue, we applied FastGxC to multi-tissue RNA-Seq data from the GTEx consortium (N = 698 individuals, 49 tissues) [12], identifying cis-eQTLs and assessing their tissue specificity. To assess performance in single-cell data, we applied FastGxC to peripheral blood mononuclear cells (PBMCs) from the CLUES [14] and OneK1K [15] cohorts (N = 237 and N = 981 individuals, respectively, across 8 cell types), and performed meta-analysis to map cis-eQTLs and evaluate their cell type specificity (Online Methods). Before quantifying cis-regulation, we confirmed that FastGxC reduces background noise, as the top principal components (PCs) of the decomposed expression data showed minimal correlation with technical and biological covariates compared to the original GTEx data (Figure S1).

We then assessed the extent of cis regulation and how context-specific these effects are across tissues and cell types (Figure 3). We identified a total of 24,196 eGenes across tissues (70.21% of tested genes) and 4,564 eGenes across cell types (29.05% of tested genes), defined as genes with at least one eQTL in any context (hierarchical FDR (hFDR) ≤ 5%, Table S5). The majority of FastGxC eGenes (86.5% and 82.7% across tissues and cell types) had at least one shared eQTL (Figure 3A), aligning with previous observations of widespread cis regulation and eQTL sharing [12, 34]. Despite extensive sharing, effect sizes varied substantially between contexts, with 72.1% of tissue eGenes and 63.9% of PBMC cell-type eGenes harboring at least one context-specific eQTL (Figure 3A). Notably, most of these context-specific eQTLs overlapped with shared eQTL loci (81.3% and 73.1% across tissues and cell types), suggesting that context-specificity often arises from effect size heterogeneity rather than the presence of an eQTL in a single context (Figure 3A). Representative examples illustrating shared-only and shared-plus-specific effects are discussed in the supplement (Figure S19).

**Figure 3.**
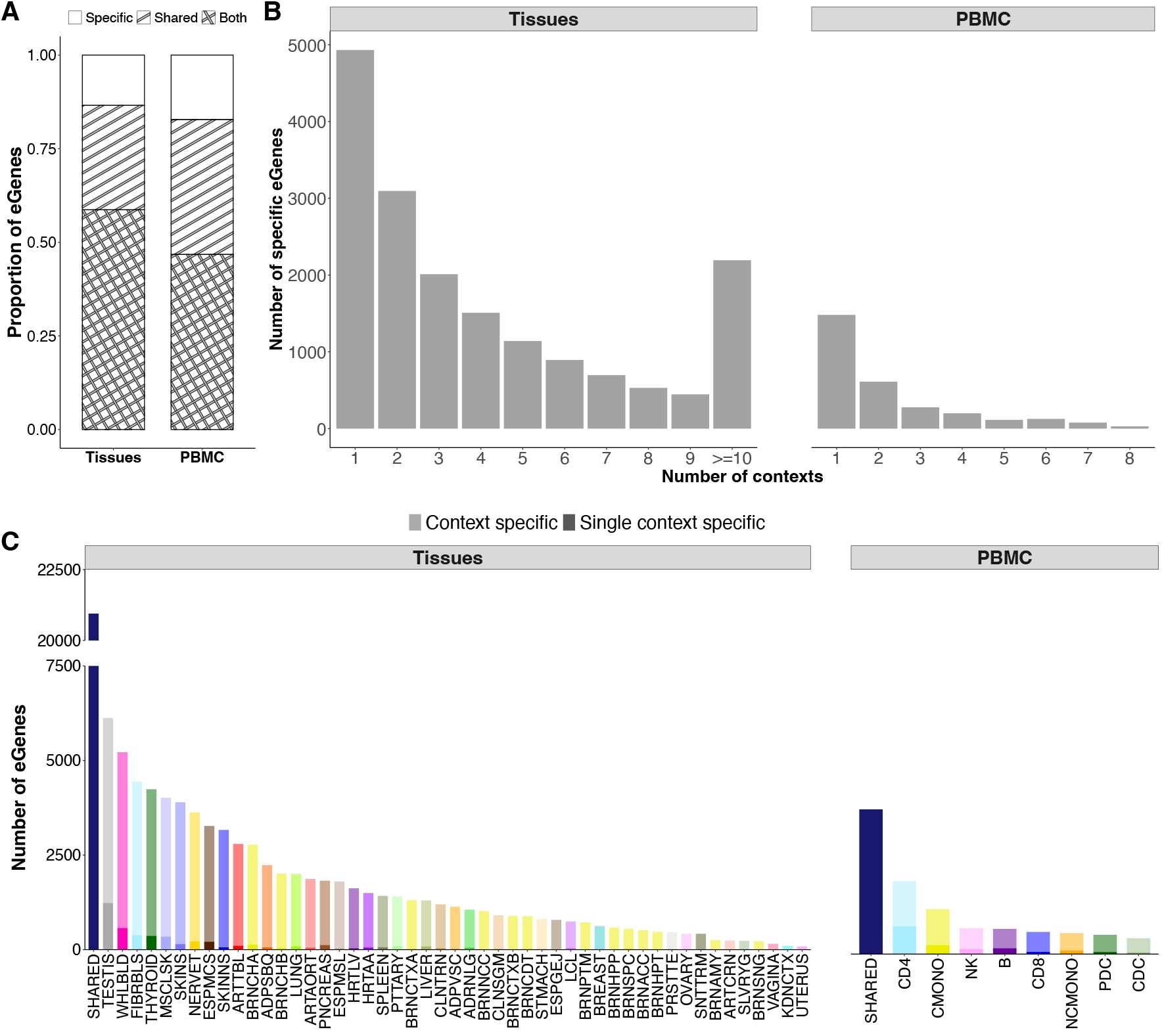
Context-specific eQTL mapping in bulk tissues and single-cell PBMC cell types. **A.**Percent of eGenes with shared-only (“Shared”), specific-only (“Specific”), and both specific and shared (“Both”) eQTLs across all tissues (left) and PBMC cell types (right). **B**. Number of contexts that drive the effect size heterogeneity for eGenes with context-specific eQTLs across tissues (left) and PBMC cell types (right). **C**. Number of eGenes with shared and context-specific eQTLs per context. For eGenes with context-specific eQTLs, opacity of color indicates the number of eGenes with specific eQTLs shared with other contexts (lightest opacity) or specific eQTLs unique to that context (darkest opacity). Tissue and cell type abbreviations are explained in Table S3.

We next aimed to determine how many and which context(s) drive the effect size heterogeneity for eGenes with context-specific eQTLs (Figure 3B-C and S3. In both bulk and single-cell data, we observed that the majority of specific eQTLs are identified in only a few contexts (Figure 3C). In tissues, much of the heterogeneity is driven by testis (6,124 eGenes), followed by whole blood (5,219 eGenes), consistent with findings from studies mapping eQTLs specific to a single tissue. [12]. Testis, which is biologically distinct from other GTEx tissues, also contributes the largest proportion (16%) of single-context-specific eQTLs—i.e., eQTLs unique to a single tissue—highlighting FastGxC’s ability to detect biologically meaningful context-specific regulation (Figure 3D). For PBMC cell types, CD4 cells (1,907 eGenes) and classical monocytes (980 eGenes) account for the majority of the heterogeneity and also contain the highest number of eQTLs unique to a single cell type. Both the number of specific eGenes and single-context specific eGenes per context are strongly correlated to the number of samples per tissue and the number of cells per cell type (Figure S18A), indicating that we may not yet have reached saturation in identifying these context-specific regulatory effects. We next compared the eGenes identified by FastGxC to those identified by the CxC approach. Consistent with our simulation results, FastGxC identified substantially more eGenes than CxC, detecting an additional 2,159 eGenes in bulk tissues and 679 eGenes in PBMC cell types (Figure S18B). Broadly, most (96.6% in tissues and 62.8% in PBMCs) FastGxC shared eQTLs overlapped with eQTLs detected in multiple contexts by CxC (Figure 4A). Importantly, FastGxC single-context-specific eQTLs mapped almost exclusively to single-context-specific eQTLs detected by CxC, demonstrating strong concordance in these cases. However, a substantial fraction (48.1% in tissues and 43.1% in PBMC cell types) of CxC single-context-specific eQTLs corresponded to FastGxC shared-only eQTLs. This discrepancy reflects the false positive specific effects that the CxC approach tends to identify — an issue we also observed in simulation results (Figure S4). Moreover, the number of context-specific eQTLs detected by CxC showed a stronger correlation with sample size than FastGxC (Figure S18A), further highlighting its sensitivity to power differences across contexts. These results underscore the limitations of CxC approaches that define context-specificity solely by the presence of significant eQTLs in isolated contexts, rather than accounting for heterogeneity in effect sizes.

**Figure 4.**
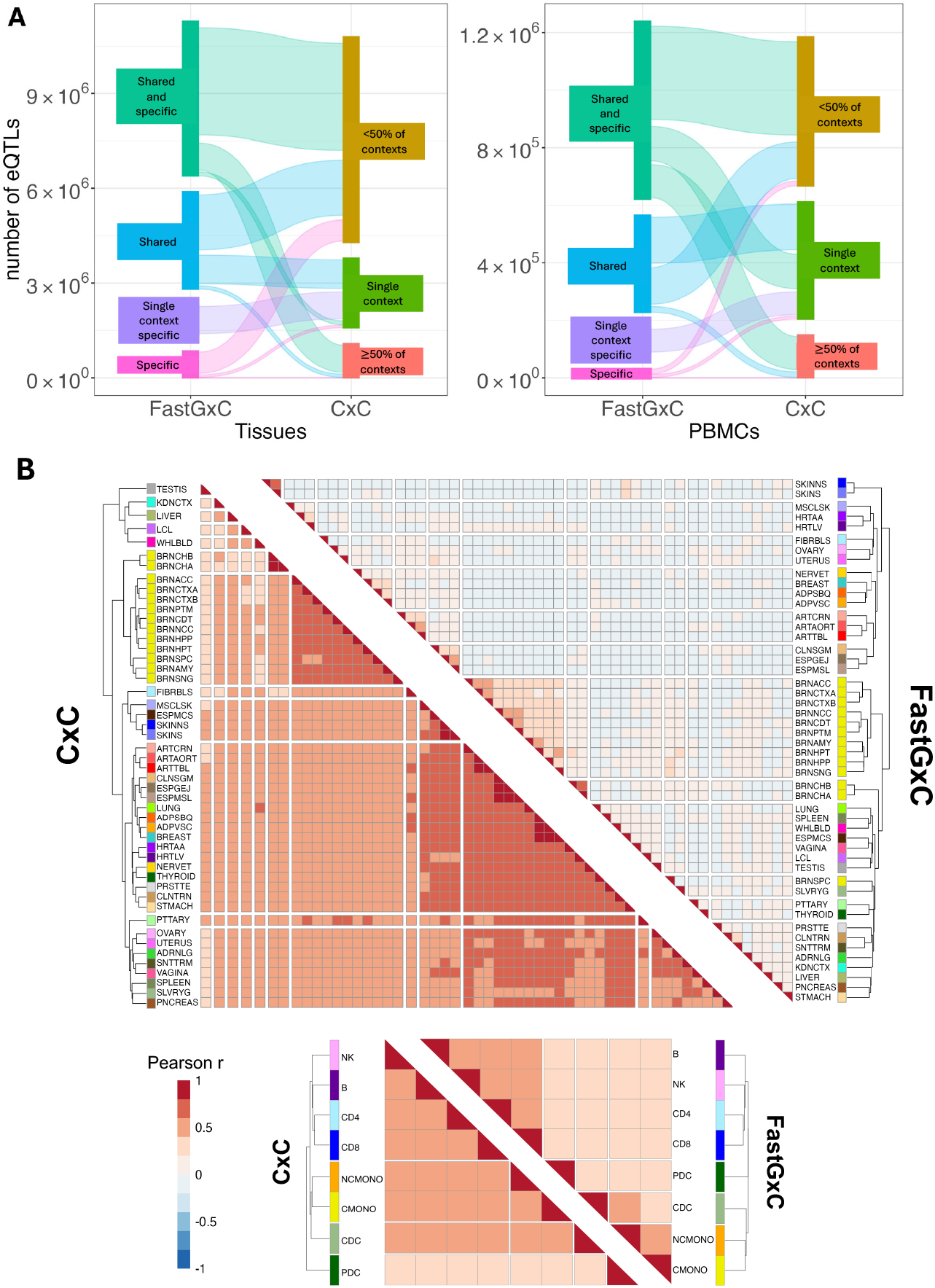
FastGxC specific eQTLs are concordant with CxC, and have strong effect size correlations among biologically related contexts. **A.**Sankey diagrams showing how eQTLs identified by FastGxC match to eQTLs identified by CxC in both tissues (left) and PBMCs (right). Node colors represent categories of eQTLs classified by CxC and FastGxC. FastGxC categories include single-context-specific eQTLs (single context specific), eQTLs that are shared or specific only (Shared, Specific), and eQTLs that are both shared and specific (Shared and specific). CxC categories include eQTLs that are found only in a single context (single context), eQTLs found in more than 1 context but *<*50% of contexts (*<*50% of contexts), and eQTLs found in ≥50% of contexts (≥50% of contexts). **B**. Heatmap with Pearson’s correlation of CxC eQTL effect sizes (left) and FastGxC context-specific eQTL effect sizes (right) across tissues (top) and PBMC cell types (bottom). Tissue and cell type abbreviations are explained in Table S3.

Finally, we show that context-specific eQTL effect sizes are correlated within groups of biologically related tissues and cell types. For example, we see that context-specific eQTL effects are correlated among 13 brain, two heart (left ventricular and atrial appendage), two artery (tibial and aorta), two esophagus (muscularis and gastro-esophageal junction), three adipose (visceral, subcutaneous, and breast), and two intestine tissues (Figure 4B - right triangle). In addition, context-specific eQTL effects are correlated most across CD4 and CD8 cells, NK cells, and B cells, between plasmacytoid and conventional dendritic cells, as well as between classical and non-classical monocytes. Furthermore, while FastGxC context-specific eQTL effect sizes show little to no correlation outside groups of biologically related tissues and cell types, CxC effect sizes show widespread correlation across all tissues and cell types regardless of biological relationships (Figure 4B - left triangle). This again demonstrates that FastGxC is able to disentangle tissue and cell type specific effects from shared effects.

### Context-specific eQTLs are enriched in functional genomic features from their matched context

To investigate functional differences between shared and context-specific eQTL variants, we performed enrichment analysis of regulatory genomic elements, comparing variants with only shared or only context-specific effects to MAF-matched non-eQTL variants (Figure 5A, right panel). In bulk tissues, variants with context-specific effects were enriched within enhancers (Odds Ratio [*OR*] = 1.06, *p* = 1.16×10^*—*5^, FDR≤ 5%), while those with shared effects were depleted (*OR* = 0.98, *p* = 2.87 × 10^−2^). Both sets of variants were enriched within promoters but the enrichment was stronger for variants with shared effects (*OR* = 1.14, *p* = 1.39 × 10^−37^) compared to those with specific effects (*OR* = 1.04, *p* = 3.14 × 10^−2^) only. In single cell PBMC cell types, we see a similar trend for enhancers (*OR*_*shared*_ = 0.98 and *OR*_*specific*_ = 1.02) but the enrichment is not significant after multiple testing adjustment, likely because the number of variants with only shared or specific effects is much smaller for single cell than bulk data. In addition, variants with shared effects only were enriched within promoters (*OR* = 1.10, *p* = 5.34 × 10^−13^), while those with specific effects were depleted (*OR* = 0.88, *p* = 8.26 × 10^−6^). These findings are consistent with previous observations that variants with context-specific effects are more enriched in genomic elements that confer context specificity to gene expression, such as enhancers, while variants with shared effects are more common within promoters [18, 44].

**Figure 5.**
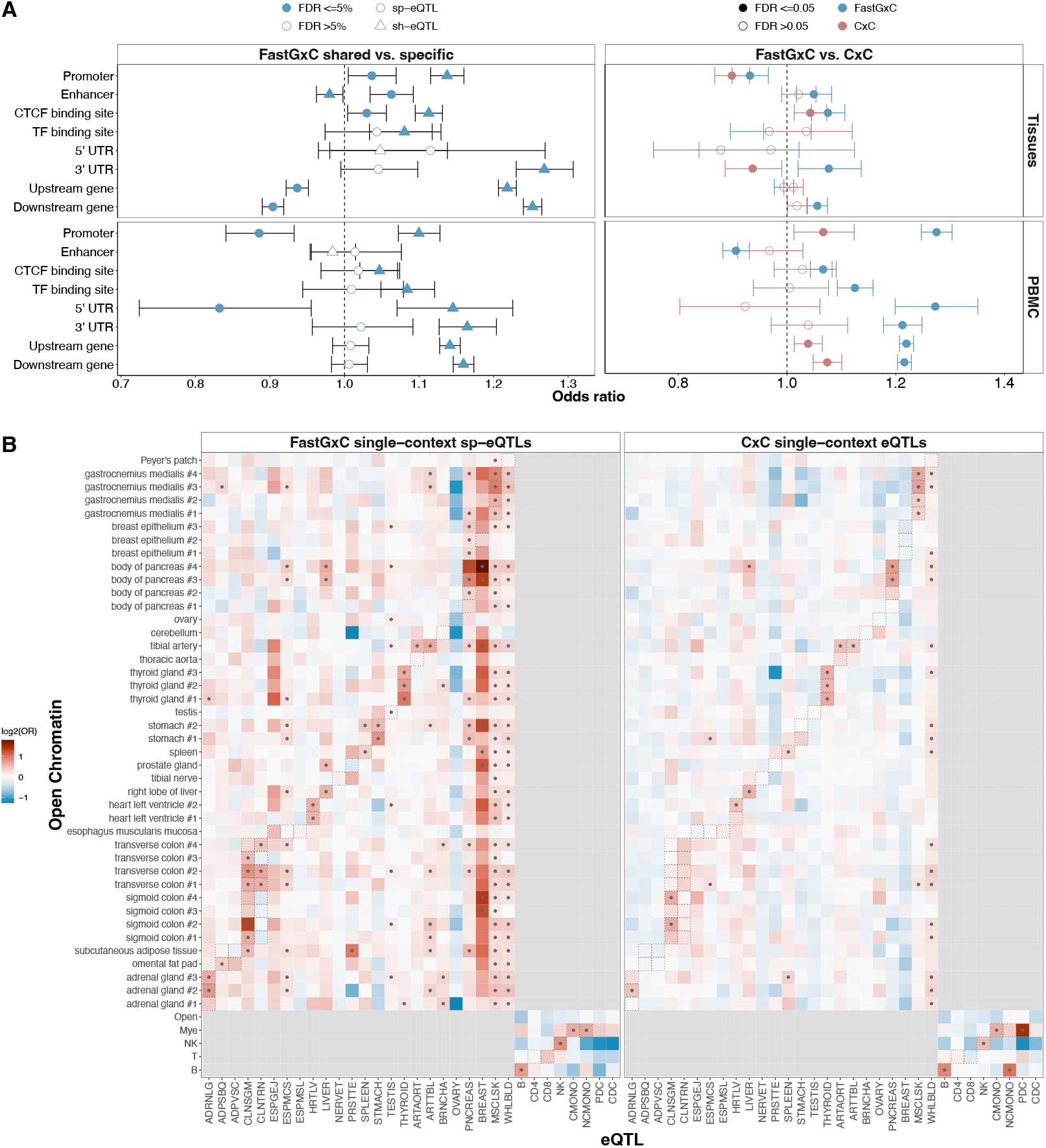
Context-specific eQTL variants are enriched in functional genomic features from their respective contexts. **A.**Enrichment of variants with FastGxC shared or context-specific effects only (left) and variants discovered by FastGxC or CxC only (right) across tissues (top) and PBMCs cell types (bottom) in genomic elements with known regulatory effects. Shape indicates different sets of variants. Color indicates different methods. Shape fill indicates significance of enrichment at *FDR* ≤ 5%. **B**. Enrichment of variants with FastGxC context-specific (left) or CxC (right) eQTL effects that are unique to a single context in regions of open chromatin across multiple tissues and cell types. Tissue and PBMC cell type open chromatin regions were obtained from ENCODE and Calderon et al. [21], respectively. Boxes indicate manual matching between chromatin and expression context. Color indicates strength of enrichment/depletion in *log*_2_ scale. Dot indicates significant enrichment at *FDR* ≤ 5%.

To understand how variants with eQTL effects mapped by the CxC approach differ functionally from those identified by FastGxC, we performed another enrichment analysis for genomic elements using sets of variants that are only discovered by CxC or FastGxC (Figure 5A right panel). Compared to CxC-only variants, FastGxC-only variants are enriched (FDR≤ 5%) in more genomic features (50% versus 12.5% of annotations in tissues and 87.5% versus 37.5% in PBMCs) and show stronger enrichment in key genomic elements, such as CTCF binding sites (*OR*_*FastGxC*_ = 1.08 and *OR*_*CxC*_ = 1.04 in tissues and *OR*_*FastGxC*_ = 1.07 and *OR*_*CxC*_ = 1.03 in PBMCs). Additionally, in tissues, FastGxC-only variants are significantly enriched in enhancers (*OR* = 1.05,*p* = 2.1 × 10^−3^), while CxC-only variants are not (*OR* = 1.02, *p* = 1.8 × 10^−1^). In PBMCs, FastGxC-only variants are depleted in enhancers but are enriched in every other genomic feature that we tested.

Chromatin is strongly context-specific [45] and therefore provides a natural framework for validating FastGxC-mapped context-specific eQTLs and quantifying the functional differences between FastGxC and CxC-mapped eQTLs. To this end, we tested for enrichment of variants with FastGxC or CxC single-context-specific eQTL effects in regions of open chromatin from matching tissues and cell types. Among bulk tissues, FastGxC variants were more often enriched in open chromatin from the corresponding tissues compared to CxC variants, with enrichment observed in 54% (29/54) versus 30% (16/54) of cases (One-sided McNemar test, p = 1.95 × 10^*—*3^; Figure 5B). Additionally, we observed widespread enrichment in open chromatin for both FastGxC and CxC variants in tissues with broadly distributed cell types, such as whole blood [46, 47]. In PBMCs, four out of six (66.6%) cell types with matching chromatin data demonstrated significant enrichment in open chromatin regions from corresponding cell types versus 50% of cell types for CxC eQTLs (One-sided McNemar test, p = 1). CD4 and CD8 cells lacked significant enrichment (FDR≤ 5%) within the broader T cell group, likely due to the aggregation of more granular subtypes, reducing specificity (see Methods) but the enrichment trend is similar (*OR*_*CD*4_ = 1.03 and *OR*_*CD*8_ = 1.34).

Together, these results highlight the functional relevance of FastGxC context-specific eQTLs, showing greater enrichment in functional genomic elements and improved capture of context-specific chromatin accessibility in matched contexts compared to CxC eQTLs. Additionally, context-specific eQTLs identified exclusively by FastGxC are more likely to reside in functional regions.

### Context-specific eQTLs identify putatively causal contexts and genes of complex traits

Mapping eQTLs is crucial for identifying the regulatory targets and context of action of disease-associated non-coding variation. To evaluate whether FastGxC eQTLs improve our understanding of the context mediating complex disease risk, we analyzed trait-associated variants from 539 traits in the NHGRI-EBI GWAS catalog [48]. Specifically, we tested for enrichment of variants with specific and shared eQTL effects identified by FastGxC in trait-associated variant sets, comparing them to an equal-sized set of MAF-matched non-eQTL variants (Table S5). Following the GTEx consortium protocol, we used expert curation to assign the most relevant tissues for each trait (Table S5) [12] and assessed precision and recall rates to identify the tissue labeled as relevant for each trait. These results were compared with those obtained from CxC eQTLs in individual contexts. PBMC cell types were excluded from this analysis due to the uncertainty regarding the exact cell type relevant for each trait.

At the same recall rate, FastGxC eQTLs achieved a three-fold increase in precision for identifying disease-relevant tissues and a two-fold improvement in their ranking compared to CxC eQTLs (Figure 6A). While CxC eQTLs typically prioritized a median of 10 out of 49 tissues per trait, likely due to widespread tissue-sharing (Figure 4B), FastGxC prioritized a median of two tissues. This suggests that modeling the extensive sharing of eQTL effects across tissues can better localize GWAS associations to a smaller, more relevant subset of tissues.

**Figure 6.**
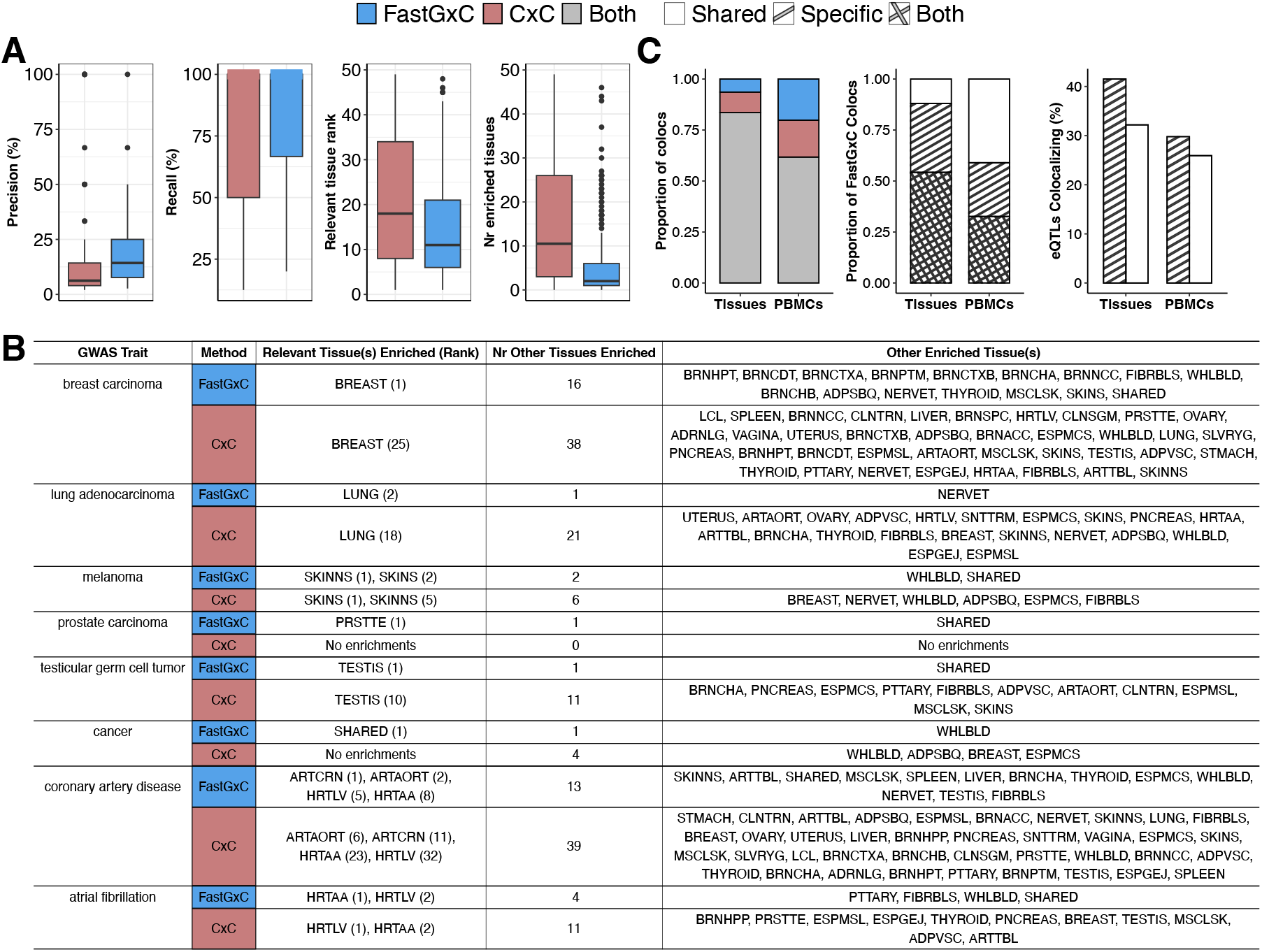
FastGxC identifies context-relevant mechanisms and increases colocalizations of complex traits. **A.**Accuracy of FastGxC and CxC eQTLs to prioritize the most relevant tissue(s) across 539 complex traits with a strong prior indication for the likely relevant tissue(s). Number of enriched tissues for each method was computed only for traits that had at least one significant enrichment in either method. **B**. Tissues prioritized by FastGxC and CxC eQTLs as well as the rank of the known relevant tissues for specific complex traits. **C**. Colocalization of FastGxC and CxC eQTLs with GWAS summary statistics across 63 human traits. **Left:** Proportion of colocalizations found uniquely by FastGxC or CxC and by both methods. **Middle:** Proportion of FastGxC identified colocalizations that are shared-eQTLs only, specific-eQTLs only, or both. **Right:** Percentage of FastGxC shared and specific eQTLs that co-localized over the total number of shared and specific eQTLs tested for colocalization for tissues and PBMCs.

Overall, FastGxC enrichment patterns aligned well with known trait-tissue associations (Figure 6B, FDR ≤ 5%). In cancer traits, where the relevant tissue is typically well-defined, FastGxC demonstrated superior tissue localization compared to CxC. For instance, in breast carcinoma, FastGxC showed the strongest enrichment in breast mammary tissue (*OR* = 5.0, *p* = 3.2 × 10^−4^), while CxC prioritized EBV-transformed lymphocytes, with breast mammary tissue ranking 25th (*OR* = 2.24, *p* = 7.5 × 10^−4^). In lung adenocarcinoma, CxC identified significant associations in 22 tissues, many unrelated to lung physiology (lung *OR* = 2.83, ranked 18th, *p* = 1.6 × 10^−3^), whereas FastGxC found associations only in lung (*OR* = 5.67, *p* = 2.60 × 10^−3^) and nerve tibial (*OR* = 20, *p* = 2.1 × 10^−5^). For traits not specific to a single tissue, such as the “any cancers” trait, FastGxC showed the strongest enrichment for shared eQTLs, consistent with processes common across tissues. This improved tissue resolution was also evident in non-cancer traits. For example, in coronary artery disease, FastGxC identified significant associations in 17 tissues, compared to 43 for CxC, with the top tissues being cardiovascular-relevant, such as coronary (*OR* = 13.0, ranked 1st) and aortic (*OR* = 2.96, ranked 2nd) artery, heart left ventricle (*OR* = 2.82, ranked 5th), and atrial appendage (*OR* = 2.37, ranked 8th).

To evaluate the ability of FastGxC eQTLs to identify the regulatory targets of trait-associated variants, we performed a colocalization analysis integrating GWAS summary statistics for 63 complex traits and diseases with FastGxC shared and specific eQTLs in bulk tissues and single-cell PBMC types (Figure 6C and Table S6). We compared these results to colocalizations based on CxC eQTLs mapped separately in each tissue and PBMC cell type. Across all traits and methods, we prioritized candidate causal genes for 5,726 (47.12% of tested) GWAS loci at a colocalization posterior probability (CLPP) threshold of 50%. The majority of the colocalizations (83.56% in tissues and 61.75% in PBMC cell types) were identified by both methods, while 6.40% and 20.18% were unique to FastGxC in tissues and PBMC cell types, respectively. This represents a 6.84% and 25.28% increase in significant colocalizations for tissues and PBMCs, respectively (Figure 6C), with the percentage increase remaining relatively consistent across CLPP thresholds for tissues and reaching up to 50% for PBMCs (Figure S20).

Previous studies suggest that context-specific eQTLs are more enriched for disease associations than shared eQTLs ([12, 14]). To test this hypothesis, we compared the colocalization rates of FastGxC shared and specific eQTLs. In tissues, most colocalizations (54.25%) involved eQTLs with both shared and context-specific effects, while 33.80% were specific-only eQTLs. In PBMCs, colocalizations were highest for shared-only eQTLs (40.93%), followed by those with both shared and specific effects (32.62%, Figure 6C). However, after normalizing by the number of shared and specific eQTLs tested for colocalization, specific eQTLs showed higher colocalization rates than shared eQTLs in both tissues (41.52% vs. 32.19%, One-sided Binominal proportion test *p* = 1.25 × 10^*—*91^) and PBMCs (29.78% vs. 25.93% *p* = 0.108) (Figure 6C). This represents a 1.2-fold enrichment in the ability of specific eQTLs to identify candidate causal genes for trait-associated variants, reinforcing their disease relevance.

Taken together, we demonstrate that FastGxC-specific eQTLs enhance the resolution of context-trait associations, increase the number of candidate causal genes for human traits, and are more disease relevant than shared eQTLs.

## 3 Discussion

We developed FastGxC, a novel statistical method for efficiently and powerfully mapping context-specific eQTLs by leveraging the correlation structure of functional genomic studies with repeated sampling. Through simulations, we demonstrated that FastGxC is well-calibrated for both identifying eQTLs and assessing their context specificity. Furthermore, FastGxC provides unbiased estimates of overall eQTL effect sizes in each context, with only slight bias in cases of extensive missing data (over 63% of data missing). FastGxC matches the power of LMM-GxC—the only other properly calibrated method for context-specific eQTL mapping—while being orders of magnitude faster.

We applied FastGxC to bulk multi-tissue and single-cell RNA-seq data sets and identified 17,447 tissue-specific and 2,920 cell-type-specific eGenes. The majority of context-specific effects appeared in loci that exhibited context-shared effects, highlighting the importance of defining context-specificity by effect size heterogeneity rather than the presence or absence of significant eQTL effects in each context. In addition, we found that context-specific eQTLs are shared mostly between groups of biologically related contexts and are more frequently enriched in genomic elements that confer context specificity to gene expression, e.g., enhancers and context-specific regions of open chromatin, providing further evidence of their validity. Finally, we found that context-specific eQTLs provide increased precision for identifying disease-relevant contexts compared to CxC eQTLs, and FastGxC specific eQTLs provided a 1.2 fold increase over shared eQTLs to identify putative causal genes that drive human traits, confirming their utility in understanding the regulatory mechanisms underlying complex human diseases.

Despite its advantages, FastGxC has certain limitations that warrant consideration. For single-cell RNA-seq data, FastGxC operates on pseudo-bulked data, aggregating expression profiles across cells within the same context. While this approach may lead to power loss in cases of substantial cell-to-cell heterogeneity within cell types, prior studies suggest that pseudo-bulk methods perform comparably to single-cell approaches [31, 49]. Additionally, FastGxC relies on predefined contexts, which can be challenging in single-cell data due to the lack of a unified framework for defining and classifying cell types [50]. Finally, while FastGxC’s marginal tests are well-calibrated, their utility diminishes in cases of extensive heterogeneity, where many contexts contribute to effect size variation (Figure S14). In addition, the context driving heterogeneity can be one without a detected eQTL—e.g., if all but one context have a significant eQTL, the remaining context will exhibit the largest (absolute) context-specific effect size (Figure S14). This highlights the nuances of interpreting context specificity in these scenarios. Nevertheless, in real-world data, FastGxC performs well, as evidenced by the enrichment of its context-specific eQTLs in functional genomic annotations and disease associations.

Additional extensions of FastGxC have the potential to further improve the power and scalability of the method, but we leave these directions to future work. The current implementation models a single shared component across all contexts, which performs well in many datasets. However, this formulation does not identify which specific contexts contribute to the shared signal and may fail to capture finer subgroup structures, such as sets of closely related tissues (e.g., brain regions in GTEx). We have previously shown that incorporating hierarchical decompositions can refine estimates of context-group-specific and context-specific eQTL effects [51]. Moreover, FastGxC defines specificity as deviation from the average effect across contexts, but some studies, such as time-course or environmental perturbation experiments, may require comparisons against a baseline context. Adjusting the decomposition step to accommodate these cases is straightforward and could expand FastGxC’s applicability. Furthermore, FastGxC assumes normally distributed expression residuals after rank-based inverse normal transformation. Extending FastGxC to handle non-normal phenotypes using generalized linear models [31, 33, 52] is straightforward but could be computationally costly. Similarly, FastGxC can be extended to capture non-linear genetic effects [30] but at a considerable computational cost and likely limited yield at current single cell sample sizes. Further improvements could come from integrating methods that model shared effect patterns across contexts [34] and incorporating fine-mapping approaches like [53] to refine candidate causal variants within significant loci, both of which are compatible with FastGxC.

In conclusion, we show that accounting for the intra-individual correlation and extensive sharing of eQTLs across contexts reveals context-specific eQTLs that can aid downstream interpretation of disease-associated variants. Furthermore, we highlight the importance of defining context specificity based on effect size heterogeneity, rather than relying on heuristic definitions and miscal-ibrated tests. We anticipate that applying FastGxC to the growing number of multi-context bulk and single-cell RNA-Seq studies will significantly expand our understanding of the context-specific gene regulatory mechanisms underlying complex human diseases.

## Supporting information

FastGxC_sup

## Acknowledgements

B.B. is supported by NIH grants U01HG012079, R01MH125252, R01DK132775, and R01DK137889. M.T. is supported by EMBO Fellowship ALTF 266-2023. L.K. is supported by NIH T32 781460-VA-32187-05. I.C is supported by a Ikerbasque Research Fellowship, funded by the EU (H2020-MSCA-COFUND-2020-101034228-WOLFRAM2), and grant PID2023-148986OB-I00 funded by MCIU/AEI/FEDER-EU. N.Z. is supported by NIH grants R01HG006399, R01CA227237, R01CA227-466, R01ES029929, R01MH122688, U01HG009080, R01HG011345, R35GM133531, R01HL155024, R01MH125252, DoD grant W81XWH-16-2-0018, and the Chan Zuckerberg Science Initiative. C.J.Y. is supported by the NIH grants R01AR071522, R01AI136972, R01HG011239, and the Chan Zuckerberg Science Initiative, and is an investigator at the Chan Zuckerberg Biohub and a member of the Parker Institute for Cancer Immunotherapy. J.W.K. is suppored by the NIH grants P30 DK116074 (to the Stanford Diabetes Research Center), R01 DK116750, R01 DK120565, R01 DK106236, R01 DK139112. The GTEx Project was supported by the Common Fund of the Office of the Director of the National Institutes of Health, and by NCI, NHGRI, NHLBI, NIDA, NIMH, and NINDS.

## Author contributions

B.B. conceived of the project and developed the statistical methods. A.L., C.L., and B.B. implemented the comparisons with simulated data. A.L., L.K., B.B., M.T., A.R., and M.G.G. performed the analyses of the GTEx, OneK1K, and CLUES data and additional analyses. A.R. and L.K. performed the colocalization analysis. B.B., A.L., and L.K. implemented the software. A.L., L.K., and B.B. wrote the manuscript, with significant input from N.Z., C.J.Y., A.D., M.G.G., and M.T. A.L., L.K., and B.B. prepared the online code and data resources. All authors read and approved the manuscript.

## Data and code availability

The FastGxC software is available as an R package at https://github.com/BalliuLab/FastGxC. All data and code to reproduce the manuscript Figures are available at https://github.com/BalliuLab/FastGxC_Manuscript. The map of shared and context-specific eGenes, as well as GWAS enrichment and colocalization results for all GTEx tissues and all PBMCs, are available as supplementary tables - S6. Data used to replicate GTEx and PBMC eQTLs are available via the GTEx portal and GEO (see Online Methods).

## Online Methods

### Overview of FastGxC method

Let *E*_*ic*_ be the observed expression of a gene for individual *i* (*i* = 1, …, *I*) in context *c* (*c* = 1, …, *C*). FastGxC first decomposes *E*_*ic*_ into an offset term, a context-shared component, and a context-specific component [54], i.e.

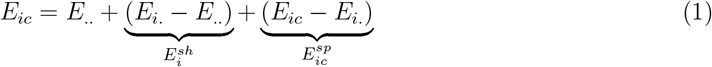

where 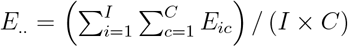 is the average expression of the gene, computed over all *I* individuals and all *C* contexts, and 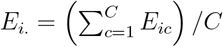 is the average expression of the gene for individual *i*, computed over all contexts. In (1), *E*_.._ is a term that is constant across individuals and contexts for each gene, 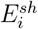 is the context-shared expression component for individual *i* and is constant across contexts for each gene and individual, and 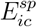 is the context-*c*-specific expression component for individual *i*.

Next, FastGxC estimates one shared and *C* context-specific cis genetic effects by regressing the genotypes on each component using ultra-fast implementations of fixed-effect linear regression models [38], i.e.,

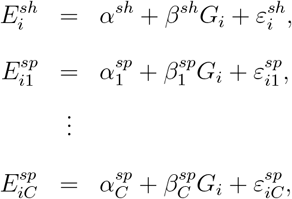

where 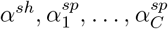 are intercepts. *G*_*i*_ ∈ {0, 1, 2} is the genotype of individual *i*, coded as number of minor alleles, and 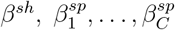 are the genetic effects on the shared and each of the context-specific expression components. Finally, 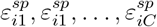 are each normally distributed residual errors with mean zero and variances 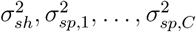.

Finally, to account for multiple testing across genes, SNPs, and contexts, FastGxC employs the hierarchical FDR-controlling procedure implemented in [42] (Figure S2). We define a gene-SNP pair as an eQTL if the SNP has a significant effect on the shared or any of the specific components of gene expression, i.e., if the global null hypothesis 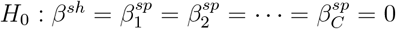 is rejected. If an eQTL exists, we define a *shared-eQTL* as a variant with a statistically significant effect on the shared expression component, i.e. if *H*_0_ : *β*^*sh*^ = 0 is rejected, and a *context-specific eQTL* as a variant with a statistically significant genetic effect on at least one context-specific expression components, i.e., if the global null hypothesis 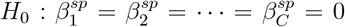 is rejected (Figure S2). In addition, we define a *specific-eQTL in context c* as a variant with a statistically significant genetic effect on the context-*c*-specific expression component, i.e., if the marginal null hypothesis 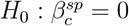 is rejected.

### Relationship between FastGxC, CxC, and LM(M)-GxC parameters

FastGxC’s eQTL effect estimates can be viewed as a computationally efficient reparametrization of those obtained through CxC and LMM-GxC approaches. Specifically, let *β*_*c*_ represent the eQTL effect in context *c*, estimated by fitting a linear regression model for each context, i.e., *E*_*ic*_ = *α*_*c*_ + *β*_*c*_*G*_*i*_ + *ε*_*ic*_. Then, the CxC eQTL effect in context *c* is equal to the sum of the shared and context-c-specific eQTL effects from FastGxC, i.e. 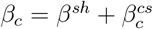. In addition, let *β*_1_ be the eQTL effect in an arbitrarily defined reference context and δ_*c*_ be the interaction eQTL effects for the non-reference context *c* from an L(M)M model with a genotype-by-context interaction term, i.e. 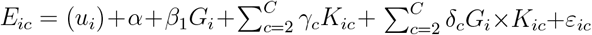. Then, 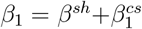 and 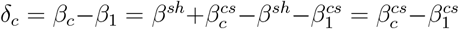. Full details of the analytical derivation are provided in the Supplementary Text.

### Simulation study

Genotypes were simulated using a binomial distribution with a minor allele frequency of 0.2. Gene expression data were generated under 35 scenarios, varying intra-individual correlation from 0 (independent contexts) to 0.8 and the cis-variant effect in each context (Table S2). Under the null hypothesis of no context-specific eQTLs (No heterogeneity), the eQTL effect was either absent across all contexts (No shared eQTL) or identical across contexts (Shared eQTL), with effect sizes explaining 5% of gene expression variability, consistent with prior estimates of cis-genetic contributions to gene expression heritability [55, 56].

Under the alternative hypothesis of eQTL effect size heterogeneity, we simulated three scenarios: (i) Single-context heterogeneity, where the eQTL explained 5% of variability in one context and 0% in others (No shared) or 10% in one context and 5% in others (Shared); (ii) Two-context heterogeneity, using similar effect size patterns; and (iii) Extensive heterogeneity, where effect sizes varied across all contexts, ranging from 0% to 10%. For each scenario, we simulated 1,000 datasets. To assess the impact of sample size, number of contexts, and missing data, we varied the number of individuals (100 or 698), contexts (8 or 49), and the proportion of missing expression data (approx. 63% and 7% across individuals and contexts), reflecting patterns observed in the GTEx [12] and OneK1K [15] data.

We obtained global estimates of type I error rates and power to identify an eQTL and test whether the eQTL was context-specific as follows. For the CxC-based approach, we used the MatrixEQTL R package [57] to fit linear regression models for the effect of the eQTL on expression in each context *c*, i.e., *E*_*ic*_ = *α*_*c*_ + *β*_*c*_*G*_*i*_ + *ε*_*ic*_, and obtained *t*-test p-values for the null hypothesis of no eQTL effect in context *c, H*_0_ : *β*_*c*_ = 0. Following the hierarchical FDR-controlling procedure implemented in [42], we then tested the global null hypothesis of no eQTL effect across contexts, *H*_0_ : *β*_1_ = … = *β*_*c*_ = 0, using Simes’s method [58], as implemented in the mppa R package [59], to combine the *t*-test p-values. Global Type I error rate and power to identify an eQTL were computed as the proportion of datasets in which the eQTL Simes’ p-value was significant at the *α* = 5% level. For the MetaTissue approach, we followed the procedure implemented in [43] and obtained the RE2 p-values which assume no heterogeneity under the null to test the null hypothesis of no eQTL effect. Global Type I error rate and power to identify a single-context-specific eQTL were computed as the proportion of datasets in which the *t*-test p-value was significant in only one context at *FDR <* 5% for CxC and M-value *>*0.9 in exactly one context for MetaTissue.

We used a similar strategy for FastGxC. Specifically, we fitted linear regression models for the effect of the eQTL on the shared and each of the *C* specific components of expression *c*, i.e., 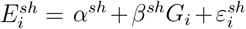 and 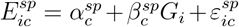, and obtained *t*-test p-values for the null hypothesis of no shared or context-*c*-specific eQTL effect, i.e., *H*_0_ : *β*^*sh*^ = 0 and *H*_0_ : *β*_*c*_ = 0. We then tested the global null hypothesis of no eQTL effect across contexts, 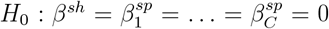, and the global null hypothesis of no context-specific eQTL effect in any context, 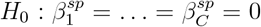, using Simes’ method to combine the corresponding p-values (Figure S2). We computed the global Type I error rate and power as the proportion of datasets in which the eQTL Simes’ p-value was significant at the *α* = 5% level.

Finally, for the LM-GxC approach, we fitted one linear model with a genotype-by-context interaction term 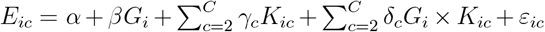 and tested the null hypothesis of no eQTL (*H*_0_ : *β* = δ_2_ = … = δ_*C*_ = 0) as well as the null hypothesis of no context-specific eQTL (*H*_0_ : δ_2_ = … = δ_*C*_ = 0) using likelihood ratio tests (LRT). For the LMM-GxC approach, we fitted one linear random effects model with a genotype-by-context interaction term 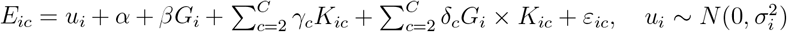 using the `lme4 R` package [60] and tested the same null hypotheses as the LM-GxC model.

To assess the ability of FastGxC to identify the heterogeneous context(s), we also obtain marginal estimates of type I error rate and power within each context by testing *C* null hypotheses of no context-specific eQTL in each context 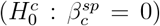 using the *t*-test implemented in the `MatrixEQTL R` package [38] and adjust for multiple testing across contexts using the Benjamini-Hochberg procedure [61]. To compare FastGxC estimates with true simulated effect sizes, we report the mean effect size across 1,000 simulated datasets for each scenario with a 95% confidence interval computed as 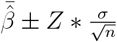 where *Z* is the z-score at *α* = 0.05.

### GTEx data quality control

Fully processed, filtered, and normalized gene expression matrices (in BED format) as well as meta data including genotype PCs, PEER factors, etc (see GTEx paper [12] for more details) for each tissue across 698 individuals were downloaded through the GTEx portal (https://www.gtexportal.org/home/datasets) on March 11, 2020. Only genes expressed (as defined by the GTEx consortium [12]) in at least two tissues were considered for downstream analyses. Prior to eQTL mapping, gene expression matrices were residualized for major sources of expression variability, including PEER factors, as per the GTEx Consortium[12]).

WGS genotype VCF data were downloaded from dbGap (dbGaP Accession phs000424.v8.p2), and only individuals with both genotype and gene expression data were retained (N=698). The VCF files were processed using vcftools (v0.1.16) to retain only bi-allelic SNPs, with variants filtered to include only those with minor allele frequencies greater than 5% in the tissue of interest. Genotype files were annotated with rs IDs using bcftools (v1.12) [62], and Plink (v1.90) [63] was used to transpose and convert the VCF files into a sample-by-genotype matrix, which served as input for eQTL mapping.

### OneK1K Genotype QC and Imputation

Array genotype data for OneK1K was obtained via the Gene Expression Omnibus (GSE196830) and included genotypes for 1,104 individuals and 759,993 markers on the Illumina Infinium Global Screening Array. For downstream analyses, only individuals with gene expression data were considered (*N* = 981). Bcftools version 1.18 was used to map SNPs to the GRCh37.13 build 151 of dbSNP [64] and adjust allele strand orientation for mismatches. PLINK version 1.90 [63] was used to filter SNPs and individuals with a call rate less than 0.95 (zero individuals and 11,317 SNPs), SNPs with a Hardy-Weinberg equilibrium test p-value less than 10^*—*6^ (1,857 SNPs), minor allele frequency (MAF) below 0.01 (211,894 SNPs), and individuals with ambiguous sex labeling (one individual).

To identify ancestry outlier samples, we performed principal component analysis (PCA) jointly on the OneK1K and 1000 Genome Phase I samples. 1000 Genome Phase I data was downloaded from the EMBL-EBI public endpoint (http://ftp.ebi.ac.uk/1000g/ftp/). PCA was conducted on the merged data using PLINK version 2.0, and 3 individuals with non-European ancestry, defined as being within three standard deviations from the European mean of genetic principal components 1 and 2, were excluded. Excess autosomal heterozygosity, defined as being within three standard deviations from the mean, was computed with PLINK version 1.90 and 7 individuals were removed. A genetic relationship matrix from all autosomal SNPs was generated using KING version 2.3.1 [65]. No individuals were excluded based on a 0.125 relatedness threshold (second-degree relatives). After quality control, 499,909 autosomal SNPs from 970 individuals were retained for imputation. Imputation was performed using the Michigan Imputation Server [66] with the 1000G phase III V5 reference panel [67] and was run using Minimac4 and Eagle v2.4. For subsequent cis-eQTL analyses, 5,849,361 SNPs with imputation quality R-squared of at least 0.8 and a minor allele frequency above 0.05 were retained.

### CLUES Genotype QC and Imputation

Genotype data for 237 individuals (188 labeled as SLE and 49 labeled as Immvar) from CLUES was obtained from dbGap (accession number phs002812.v1.p1). Genotypes from the SLE and Immvar CLUES cohorts were processed and imputed separately, as in [14], resulting in 22,159,030 variants for the SLE cohort and 9,797,072 variants for the Immvar cohort. After imputation, variants with an imputation quality R-squared greater than 0.8 were retained, and the datasets were merged using PLINK v2.0, producing a combined total of 16,616,859 variants.

Based on genetically determined ancestry, individuals were then split into European (N=140) and Asian (N=97) cohorts, following the approach described in Perez et al [14]. Within each ancestry group, PLINK v2.0 was used to filter for variants with a minor allele frequency above 5%, resulting in 4,995,061 variants in the Asian cohort and 5,292,554 variants in the European cohort for further analysis. Genotype principal components were computed within each cohort using PLINK v1.90 to identify outlier individuals, defined as those within three standard deviations of the mean for genetic principal components 1 and 2. This filtering excluded 5 individuals from the Asian cohort and 8 from the European cohort.

### OneK1K and CLUES Gene Expression QC

Gene expression data for the CLUES cohort was obtained from GEO (accession number GSE174188). For downstream analysis, we retained cells from the eight cell types used for cis-eQTL analysis in [14]: B cells, conventional and plasmacytoid dendritic cells (cDC and pDC), classical and non-classical monocytes (cMono and ncMono), NK cells, CD4 T cells, and CD8 T cells. As with genotype data, the CLUES cohort was divided into European and Asian ancestry groups. Only individuals with both expression and genotypes were retained, i.e., N=140 and N=97 European and Asian ancestry individuals.

Gene expression data for the OneK1K cohort was obtained via GEO (accession number GSE196830). From the 29 pre-defined cell type clusters, we consolidated labels for similar cell types. Specifically, memory B cells, naive B cells, and transitional B cells were grouped as B cells; natural killer cells and CD16-negative, CD56-bright natural killer cells were combined as NK cells; T alpha-beta cytotoxic, CD4-positive cells, T alpha-beta, CD4-positive cells, and T regulatory cells were grouped as CD4 T cells; and T alpha-beta, CD8-positive cells, T gamma-delta cells, and T mucosal invariant cells were grouped as CD8 T cells. For downstream analyses, only cells from the eight cell types present in CLUES were retained. In addition, only individuals with both expression and genotypes were retained, i.e., N=981 individuals.

For each individual in the CLUES European, CLUES Asian, and OneK1K cohorts, a pseudo-bulk expression profile was generated by averaging counts across cells for each cell type and gene. The following steps were then applied separately within each cohort and cell type. First, expression values for each gene were adjusted for library size factors and normalized using counts per million (CPM) and transcripts per million (TPM). Genes with normalized expression greater than zero in at least 10% of samples were retained. Gene expression values were then transformed to approximate normality using the `RankNorm` function in the `RNOmni` R package [68]. Within each cohort, genes expressed in fewer than three cell types were excluded. Next, principal component analysis (PCA) was applied to identify and remove outlier samples defined as those falling outside three standard deviations from the mean of expression principal components 1 and 2. After quality control, 17,199, 16,225, and 14,701 genes and 132, 92, and 970 individuals in the CLUES European, CLUES Asian, and OneK1K cohorts, respectively, were retained for downstream analyses.

To capture major sources of expression variation, the PCA implementation in the `PCAForQTL R` package [69] was used with the `runBE` function to determine the number of principal components that explain a significant portion of variation in each cell type. Gene expression data for each cell type and cohort was then residualized for six genotype principal components, selected gene expression principal components, sex, age, batch, and, in CLUES cohorts only, SLE status.

### Expression principal components analysis and correlation with covariates

To evaluate the effectiveness of FastGxC in removing gene expression background noise, we first applied PCA separately to the original gene expression data and the decomposed shared and context-specific expression data, using the `prcomp` function in the `stats R` package. Next, we correlated technical and biological covariates with the first ten principal components (PCs) from each data. The correlation between expression PCs and covariates was computed using the `canCorPairs` function from the `variancePartition R` package ([70]). In short, when comparing two continuous variables (e.g. gPC1 or weight), Pearson correlation was used. In order to accommodate the correlation between a continuous and a categorical variable (e.g. cohort) canonical correlation analysis (CCA) was used. Note that CCA returns correlations values between 0 and 1.

### FastGxC and CxC eQTL mapping

Residualized gene expression (see methods above) for each was mean-centered across all individuals and contexts, then decomposed into 49 tissue-specific components for GTEx and 8-cell type-specific components for CLUES and OneK1K, along with one shared expression component. Cis genetic effects were estimated on shared gene expression levels (FastGxC), context-specific gene expression levels (FastGxC), and gene expression levels within each context (CxC) using ultra-fast linear regression models in the `MatrixEQTL R` package [38], with `model=modelLINEAR` and a 1 Mb window for cis-eQTL calls. The CLUES European, CLUES Asian, and OneK1K cohorts were then meta-analyzed within each cell type using METASOFT (v2.0) [71] random effect model (RE2) with default parameters.

Multiple testing correction was applied separately for CxC and FastGxC and for bulk and meta-analyzed single-cell results using the hierarchical FDR procedures in the `TreeQTL R` package [42] with an alpha level of 5% in each level. For the CxC approach, a three-level hierarchy with genes, gene-SNP pairs, and gene-SNP-context triplets was used. For FastGxC, multiple testing correction was applied separately for the shared and specific components using a two-level hierarchy for shared eQTLs (genes and gene-SNP pairs) and a three-level hierarchy for context-specific eQTLs (genes, gene-SNP pairs, and gene-SNP-context triplets) (Figure S2B).

### Correlations of eQTL effect sizes across context

Pearson correlations were computed across all tissues in GTEx using effect sizes of only the significant eQTLs in each tissue for the CxC approach or tissue-specific eQTLs for FastGxC. Missing effect size values, due to an eQTL not being tested or not reaching significance in some tissues, were set to zero, and pairwise complete correlations were calculated using the cor function in R. For single PBMC cell types, Pearson correlations of eQTL effect sizes across cell types were computed using all tested, meta-analyzed METASOFT RE2 effect sizes.

### Building set of background SNP for enrichment analyses

The `matchit` function from the `MatchIt R` package was used to create a set of background SNP-gene pairs for each variant set of interest, matched by minor allele frequency (MAF) using the nearest neighbor matching method with a 1:1 ratio [72]. Only variants used for eQTL mapping were included in building these background sets. For eQTL sets containing more than 5,000 variants, the sets were randomly split into chunks to expedite computation. This analysis was done separately in bulk-tissues and single-cell PBMCs and for FastGxC and CxC eQTLs.

### EQTL enrichment in genomic features

To test for enrichment of various eQTL types within genomic annotations, we first created variant sets specific to each type of eQTL. This analysis was conducted separately for bulk tissues and single-cell PBMC cell types. Specific-eQTL-only and shared-eQTL-only variant sets were derived by taking the set difference of specific-eQTL and shared-eQTL variants. The FastGxC eQTL variant set was constructed by combining shared- and specific-eQTL variants across contexts, and the CxC eQTL variant set was created by taking the union of eQTL variants across contexts. We then calculated the set difference between the FastGxC and CxC eQTL variant sets to obtain the final FastGxC-only and CxC-only eQTL variant sets.

Variants from each set were annotated using the Ensembl Variant Effect Predictor (VEP) tool, which identifies variant effects, such as potential impacts on protein sequences or positioning within genomic regulatory elements. Enrichment of each eQTL set within VEP-annotated categories was tested using a one-sided Fisher’s exact test from the stats R package, followed by BH correction; significance was defined as a BH-adjusted p-value less than 0.05.

### Enrichment in regions of open chromatin

For bulk tissues, all available tissue ATAC-seq data in the ‘not perturbed,’ GRCh38, and bigBed narrowPeak categories were downloaded from in November 2020. The downloaded bigBed files were converted to bed format using the UCSC `bigbedtobed` tool for downstream analysis. For single-cell PBMC cell types, cell-type-specific ATAC-seq peaks were downloaded from [21] and grouped into major cell types (B, T, NK, Myeloid, Open), following the approach of Perez et al. [14]. Bed files were then sorted using the command `bedtools sort -k1,1 -k2,2n` to enable memory-efficient processing for subsequent intersections.

Enrichment analysis of FastGxC and CxC single-context eQTL variants from both bulk tissues and single-cell PBMC cell types in regions of open chromatin was performed by intersecting each eQTL variant set of interest with each pre-sorted bed file, representing ATAC-seq peaks from a tissue, cell type, or sample (when multiple samples were available for each context), using the `bedtools intersectBed` command. A one-sided Fisher’s exact test was used to obtain the statistical significance of each enrichment, followed by BH multiple testing adjustment. Significance was called for BH-adjusted p-values less than 0.05.

### Enrichment of bulk tissue eQTLs in GWAS loci

Genome-wide association study (GWAS) data (gwas_catalog_v1.0.2-associations_e100_r2020-06-17) including 1,563 unique traits was downloaded from the NHGRI-EBI GWAS Catalog in August 2020 [48] and processed for downstream analysis. Matching of variants with and without eQTL effects was performed as previously described.

Enrichment analysis of FastGxC and CxC eQTL variants was performed by intersecting each eQTL variant set of interest with trait-associated variants from the GWAS catalog based on rs IDs. Statistical significance was assessed using a one-sided Fisher’s exact test for each enrichment. Only mapped traits within the GWAS catalog that contained at least 10 significant loci were included in our downstream analysis resulting in 539 traits with complete enrichment results. Multiple testing correction was applied using the hierarchical FDR procedures in the `TreeQTL R` package [42], with tissues at level one and tissue-trait pairs at level two, maintaining an FDR of 5% at each level.

The most likely causal tissue(s) for the 539 GWAS traits were annotated manually, following the approach used in [12]. These annotations were used to calculate precision and recall rates. Specifically, for each trait, a contingency table was constructed to capture the frequency with which a trait of interest is both enriched in a tissue’s eQTLs and assigned as the likely relevant tissue. This yielded true positive, false positive, true negative, and false negative counts (TP, FP, TN, FN). Precision was calculated as TP / (TP + FP), and recall as TP / (TP + FN).

### Colocalization of eQTLs with GWAS variants

We obtained complete GWAS summary statistics for 63 unique complex traits. The list of traits analyzed, along with their references, is available in Table S5. Colocalization analysis of GWAS variants with eQTLs from bulk tissues and single-cell PBMC cell types was conducted using a custom integration of FINEMAP [73] and eCAVIAR [74] with an LD-modified colocalization posterior probability (*CLPP*_*mod*_) following the method outlined in Gloudemans et al. [13]. Significant colocalization was defined as *CLPP*_*mod*_ above 0.5, as in [75], and impact of this threshold on results is studied in Figure S20. To assess the colocalization contribution from FastGxC specific versus shared eQTLs in PBMCs, we limited our analysis to the 10 immune related traits out of the original 63.

