## Supplementary material for "FastGxC: Fast and Powerful Context-Specific eQTL Mapping in Bulk and Single-Cell Data": FastGxC_sup

### Exact relation between FastGxC and CxC estimates

Fix a gene and assume that its expression in each context follows a linear model:

$$E_{i,}^0 = G\beta^0 + \epsilon_{i,} \in \mathbb{R}^C$$

where:

- $E^0 \in \mathbb{R}^{N\times C}$ is the matrix of gene expression across $N$ samples and $C$ contexts, e.g. tissues or cell types
- $G \in \mathbb{R}^{N\times S}$ is an arbitrary covariate matrix containing $S$ features (in this paper, the features are *cis*-SNPs, and usually $S = 1$)
- $\beta^0 \in \mathbb{R}^{S\times C}$ are the genetic effects in each context
- $\epsilon_{i,} \in \mathbb{R}^C$ captures arbitrarily distributed noise, assumed i.i.d. over samples $i$ but with covariance between contexts given by $\mathbb{V}(\epsilon_{i,}) = \Sigma$

Now define the context-centered expression as:

$$E_i = E_i^0 - \bar{E}_i 1_C^T \quad \text{or} \quad E_{ic} = E_{ic}^0 - \bar{E}_i$$

where $1_C \in \mathbb{R}^C$ is a vector of 1s and $\bar{E} \in \mathbb{R}^N$ is a vector containing each sample's mean expression across all $C$ contexts.

For any arbitrary vector $X \in R^{1\times N}$, we have:

$$XE = XE^0 - X\bar{E}1_C^T$$

In particular, when $X = \frac{1}{\|G_j\|^2}G_j$ for SNP $j$, then:

- $XE := \hat{\beta}$ gives the FastGxC cs-eQTL effect size estimates for SNP $j$
- $XE^0 := \hat{\beta}^0$ gives the ordinary eQTL effects in each contexts
- $X\bar{E} := \bar{\beta}$ gives the FastGxC sh-eQTL effects

Putting these three facts together proves:

$$\hat{\beta}_c = \hat{\beta}_c^0 - \bar{\beta} \quad \text{or} \quad \text{FastGxC} = \text{CxC} - \text{Shared}$$

for all contexts $c$. In other words, the standard context-specific estimates in CxC naturally and exactly decouple into the FastGxC estimates and the cross-context average estimate.

By the same argument, CxC decomposes into FastGxC and shared effects even when:

- Covariates are included, via $X = \frac{1}{\|P_Z^\perp G_j\|^2} G_j P_Z^\perp$, where $P_Z^\perp$ is the orthogonal projection onto the span of the covariate matrix $Z$
- Multiple SNP effects are fit simultaneously, via $X = (GG^T)^{-1}G^T$
- Ridge regression/kinship-based LMMs are used, if the regularization/heritability is equal across contexts

Conceptually, associativity guarantees that linear operators applied to the left of the matrix $E$ play well with linear operators applied to its right. And most regression involve linear operations on $E$ from the left, while the centering operation used by GxC is a linear operator from the right. That is, we can center and then perform regressions (as in FastGxC) or can perform regular regressions and then center; these operations associate, therefore give identical results.

### Approximate relation between FastGxC and CxC standard errors

Above, we showed the CxC estimates exactly decouple into FastGxC and shared estimates. Here, we show a similar result for the standard errors, though it holds only approximately. Specifically, the variance of the FastGxC estimate is roughly the variance of the CxC estimate minus the variance in the shared estimate. This provides a sharp description of the improvement in power in FastGxC over CxC due to removal of shared noise.

More concretely, using the equivalence proved above, we have:

$$
\begin{aligned}
\mathbb{V}\left(\hat{\beta}_c\right) &= \mathbb{V}\left(\hat{\beta}_c^0 - \bar{\beta}\right) \\
&= \mathbb{V}\left(\hat{\beta}_c^0\right) + \mathbb{V}\left(\bar{\beta}\right) - 2\mathrm{Cov}(\hat{\beta}_c^0, \bar{\beta}) \\
&= \mathbb{V}\left(\hat{\beta}_c^0\right) - \mathbb{V}\left(\bar{\beta}\right) - 2\left(\mathrm{Cov}(\hat{\beta}_c^0, \bar{\beta}) - \mathbb{V}\left(\bar{\beta}\right)\right) \\
&\approx \mathbb{V}\left(\hat{\beta}_c^0\right) - \mathbb{V}\left(\bar{\beta}\right) \qquad (*)
\end{aligned}
$$

Loosely, the approximation assumes that the contexts are roughly exchangeable, or that each context is roughly equally correlated with other contexts[1]. For example, this holds exactly in the cases where contexts are IID ($\Sigma = \sigma^2 I$) or exchangeable ($\Sigma = \sigma^2 I + bJ$); conversely, this is violated if context $c$ is very unique, or if there large and structured subsets of the contexts (eg brain regions).

For example, imagine that $C$ is large and that each sample's noise has exchangeable distribution across contexts, implying that $\mathbb{V}\left(\epsilon_{i,}\right) = \sigma^2 I + sJ$ for some $\sigma^2 > c$. Then the above approximation is exact, and standard error in FastGxC simply subtracts off the standard error for the shared noise term, $s$:

$$
\mathbb{V}\left(\hat{\beta}_c\right) = \frac{1}{\|X\|^2}(\sigma^2 + s) - \frac{1}{\|X\|^2}(\frac{1}{C}\sigma^2 + s) \approx \frac{1}{\|X\|^2}\sigma^2
$$

[1]More formally, if we assume that $\epsilon_i$ are i.i.d. with cross-context covariance matrix $\Sigma$, then:

$$
\mathrm{Cov}(\hat{\beta}_c^0, \bar{\beta}) = \mathrm{Cov}(XE_{,c}^0, X\bar{E}) = X\mathrm{Cov}(E_{,c}^0, \bar{E})X^T = \|X\|^2\mathrm{Cov}(E_{,c}^0, \frac{1}{C}E^0 1_C) = \frac{1}{C}\|X\|^2\Sigma_{c,}1_C = \|X\|^2\Sigma_{c.}
$$

$$
\Sigma_{c.} := \frac{1}{C}\sum_{c'}\Sigma_{cc'}
$$

and likewise (using $\otimes$ for tensor/Kronecker product, and $\mathrm{vec}(\cdot)$ for column-wise matrix vectorization):

$$
\mathbb{V}\left(\bar{\beta}\right) = \mathbb{V}\left(X\bar{E}\right) = \mathbb{V}\left(\left((\frac{1}{C}1_C^T) \otimes X\right)\mathrm{vec}(E^0)\right) = \frac{1}{C^2}\left(1_C^T \otimes X\right)\left(\Sigma \otimes I_N\right)\left(1_C^T \otimes X\right)^T = \left(\frac{1}{C^2}1_C^T\Sigma 1_C\right)\cdot(XX^T) = \|X\|^2\Sigma_{..}
$$

$$
\Sigma_{..} := \frac{1}{C^2}\sum_{c,c'}\Sigma_{cc'}
$$

Thus, $(*)$ assumes that $\Sigma_{c.} \approx \Sigma_{..}$, i.e. that context $c$ is about as correlated with the average context as any other.

### Inter-context noise correlation does not affect FastGxC estimates

Say that samples are i.i.d. Gaussian but that contexts are correlated:

$$E_{i,} \overset{\text{iid}}{\sim} G_{i,}\beta^0 + \mathcal{N}(0, \Sigma)$$

Assume that we estimated or know the noise covariance $\Sigma$, e.g. with an LMM. The GLS and OLS estimates for $B$ are identical–again, conceptually, the key fact is that column transformations on $E$ operate independently of row transformations. ($\Sigma$ acts on the rows of $E$, while $G$ acts on the columns.) One way to see this is using the covariance across all entries of $E$, $\mathbb{V}(\text{vec}(E)) = \Sigma \otimes I_N$:

$$\begin{aligned}
\hat{\beta}_{GLS} &:= \left((G \otimes I_P)^T (I_N \otimes \Sigma)^{-1} (G \otimes I_P)\right)^{-1} (G \otimes I_P)^T (I_N \otimes \Sigma)^{-1} \text{vec}(E) \\
&= \left((G^T G)^{-1} \otimes \Sigma\right) \left(G^T \otimes \Sigma^{-1}\right) \text{vec}(E) \\
&= \left(((G^T G)^{-1} G^T) \otimes I_P\right) \text{vec}(E) \\
&= \text{vec}\left((G^T G)^{-1} G^T E\right) \\
&= \hat{\beta}_{OLS}
\end{aligned}$$

**Examples of eQTLs identified by FastGxC.** To provide insight into patterns of sharing and specificity of eQTL effects revealed by FastGxC, we discuss a few individual examples (Figure S19).

First, we examine *CBS*, a gene that encodes the enzyme cystathionine beta-synthase that catalyzes the rate-limiting step of the transsulfuration pathway [76, 77] (Figure S19A). This pathway acts ubiquitously across many cell-types to perform diverse and important biological functions such as protein synthesis and methylation [78]. Indeed, eQTL effect size estimates from CxC are significant in 48 individual tissues (hFDR $\leq$ 5%), suggesting a universal, shared mechanism of genetic regulation. FastGxC crystallizes this shared mechanism by identifying a single shared-eQTL and no specific-eQTLs (hFDR $\leq$ 5%).

Second, we show an eQTL for *SIGLEC14*, an immune cell surface receptor of the immunoglobulin superfamily involved in the innate immune response [79] (Figure S19B). Similar to the *CBS* example, there seems to be a sharing of genetic effects across GTEx tissues when interpreting CxC effect sizes which could lead one to conclude that this genetic effect is invariant across the body. Yet, when we explicitly model this sharing with FastGxC, a specific-eQTL effect in whole blood emerges, indicating that, while *SIGLEC14* is under a universal tissue-shared genetic regulation, there is importantly also a blood-specific regulatory mechanism that is consistent with the known role of *SIGLEC14* in immunity.

Third, we examined the genetic regulation of *LDHC*, which encodes the testis-specific enzyme lactate dehydrogenase C, the first testis-specific enzyme discovered in male germ cells [80] (Figure S19C). We found that *LDHC* exhibits a strong positive eQTL effect in all tissues except the testis for which the eQTL effect is in the opposite direction. This lone effect becomes very apparent when specific-eQTLs are examined with FastGxC. To the best of our knowledge, this is the first time that testis-specific genetic regulation, in addition to testis-specific expression, is reported for this gene, suggesting that tissue-specificity can be regulated at multiple biological levels.

Other examples of widespread sharing obscuring a tissue-specific effect can be seen for gene *GBP3* in the spleen, gene MUC20P1 in LCLs and thyroid, and gene *GSTT2* and the two skin tissues (Figure S19D-F).

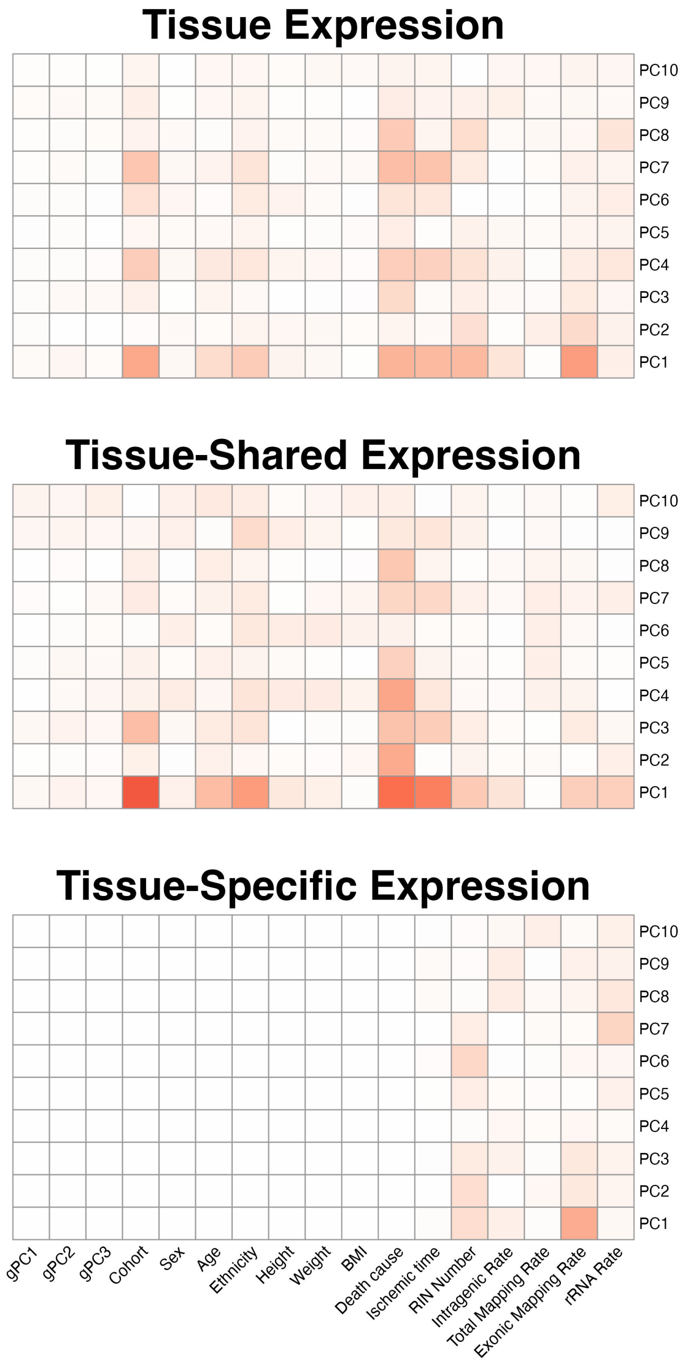


**Figure S1. FastGxC decomposition removes shared noise from context-specific expression components.** Correlation of PCs from tissue expression, tissue-shared expression, and tissue-specific expression with covariates related to study design and sample quality in GTEx. The decomposition reduces the intra-individual correlation as demonstrated by lack of correlation between PCs from the tissue-specific expression and variables that are shared/invariant within an individual across tissues, e.g. genotype PCs (gPC), sex, age, etc.

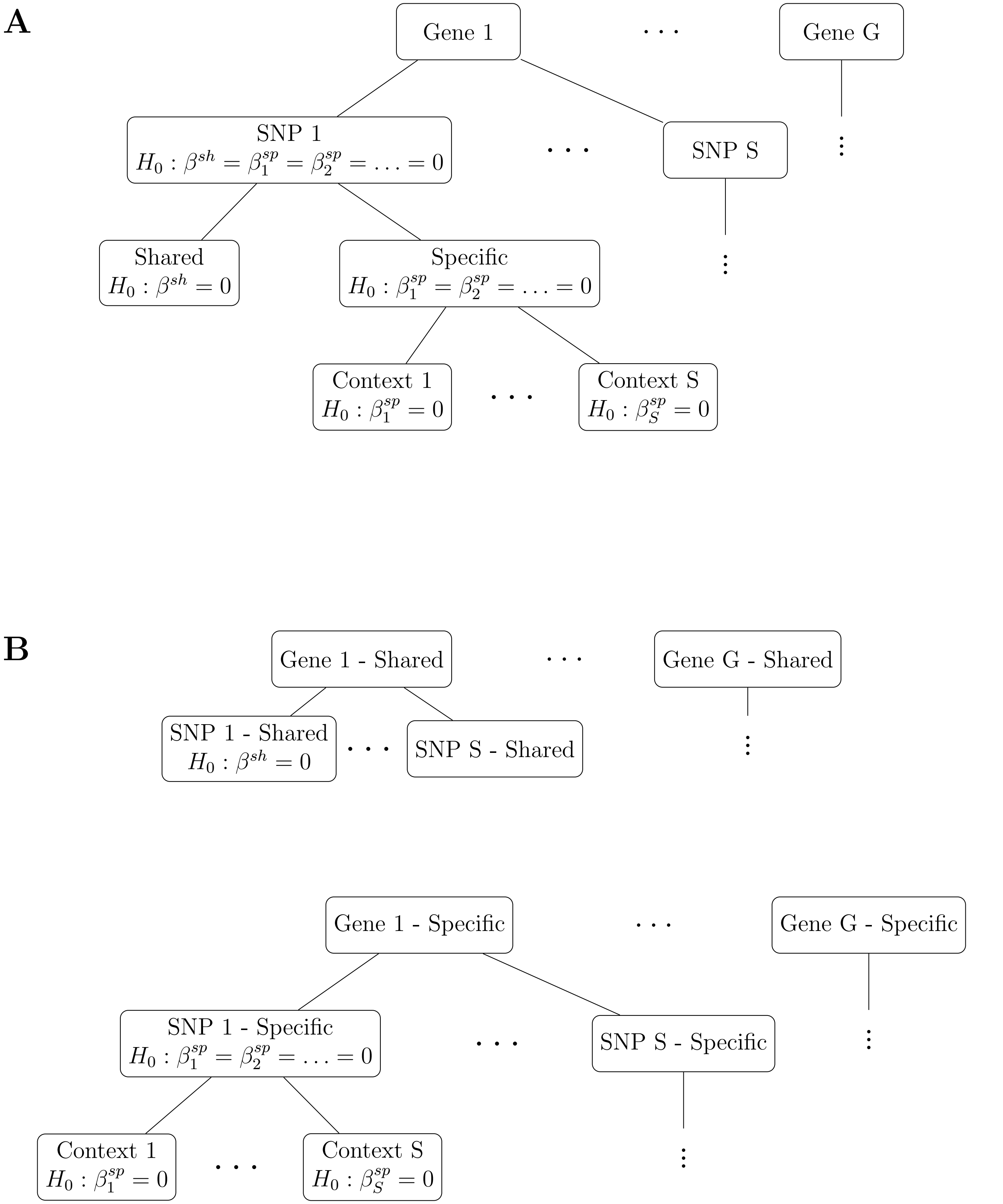


**Figure S2. Hierarchical hypotheses structures used by FastGxC. (A)** Four-level hierarchical structure used by FastGxC to adjust for multiple testing across genes (eGenes), gene-SNP pairs (eQTLs), gene-SNP-component triplets, and gene-SNP-components-contexts quadruplets. Note that in simulations, only one SNP is tested per gene so the level one and level two p-values are identical. **(B)** Two-level (top) and three-level (bottom) hierarchical structure used to adjust for multiple testing for the shared (top) and specific (bottom) eGenes / eQTLs in real data analysis.

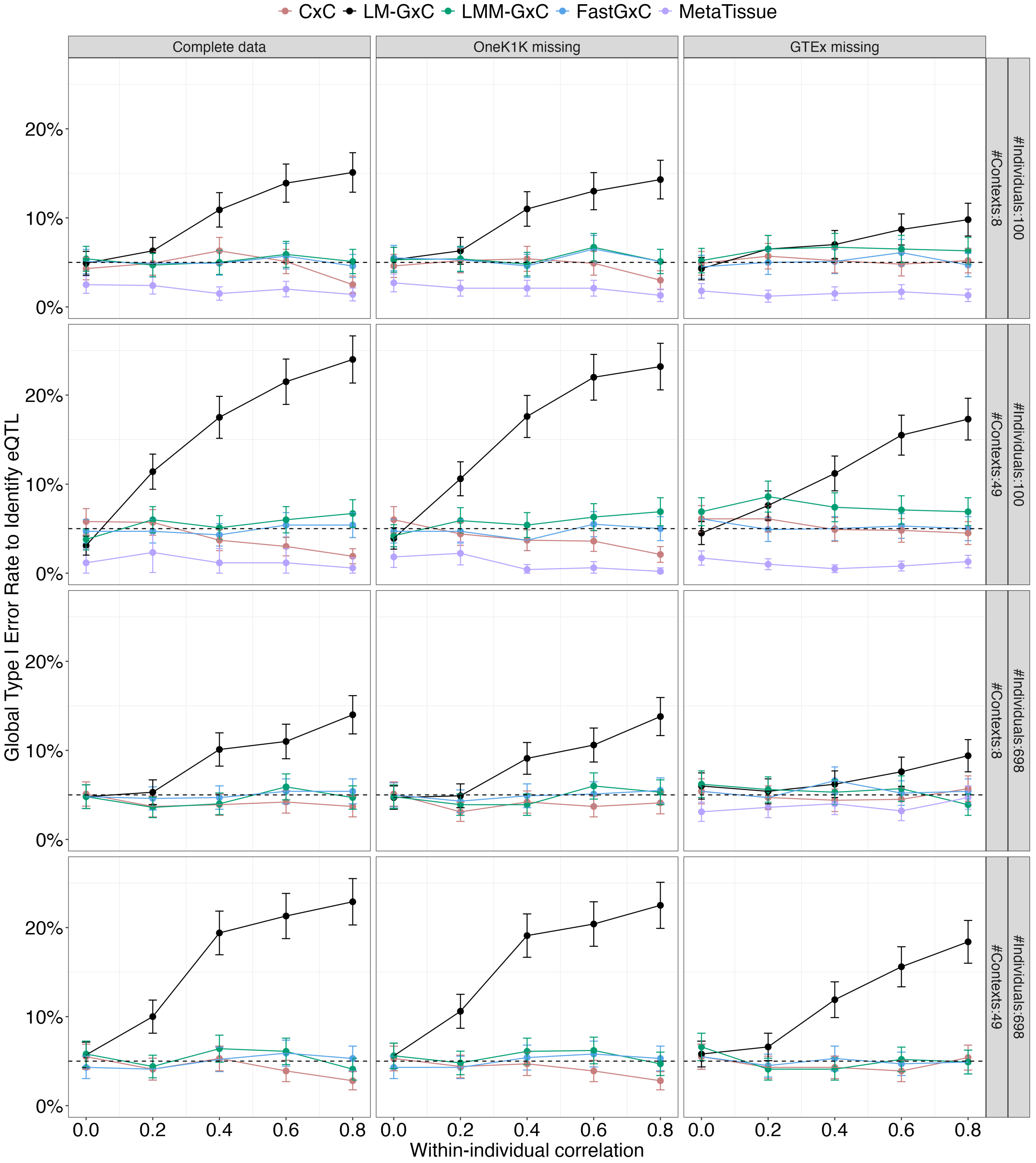


**Figure S3. Global type 1 error rate to identify an eQTL.** Global type 1 error rate (y-axis) of each method (color) to identify an eQTL as a function of within-individual residual correlation of expression (x-axis), different combinations of sample size and number of contexts (rows), and different missing data patterns(columns).

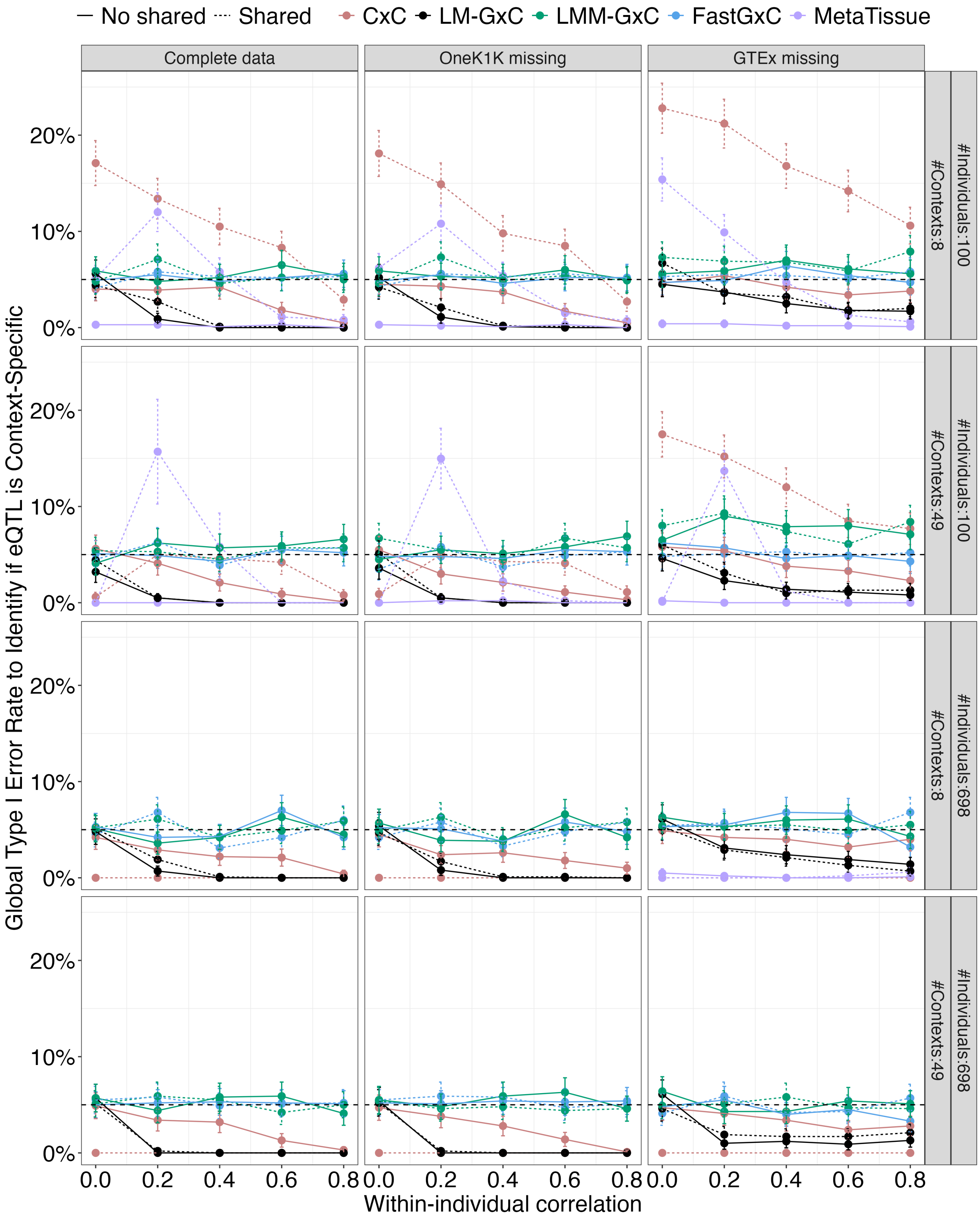


**Figure S4. Global type 1 error rate to test if eQTL is context specific.** Global type 1 error rate (y-axis) of each method (color) to test if eQTL effect is context-specific as a function of within-individual residual correlation of expression (x-axis), presence of a shared eQTL (line type), different combinations of sample size and number of contexts (rows), and different missing data patterns(columns).

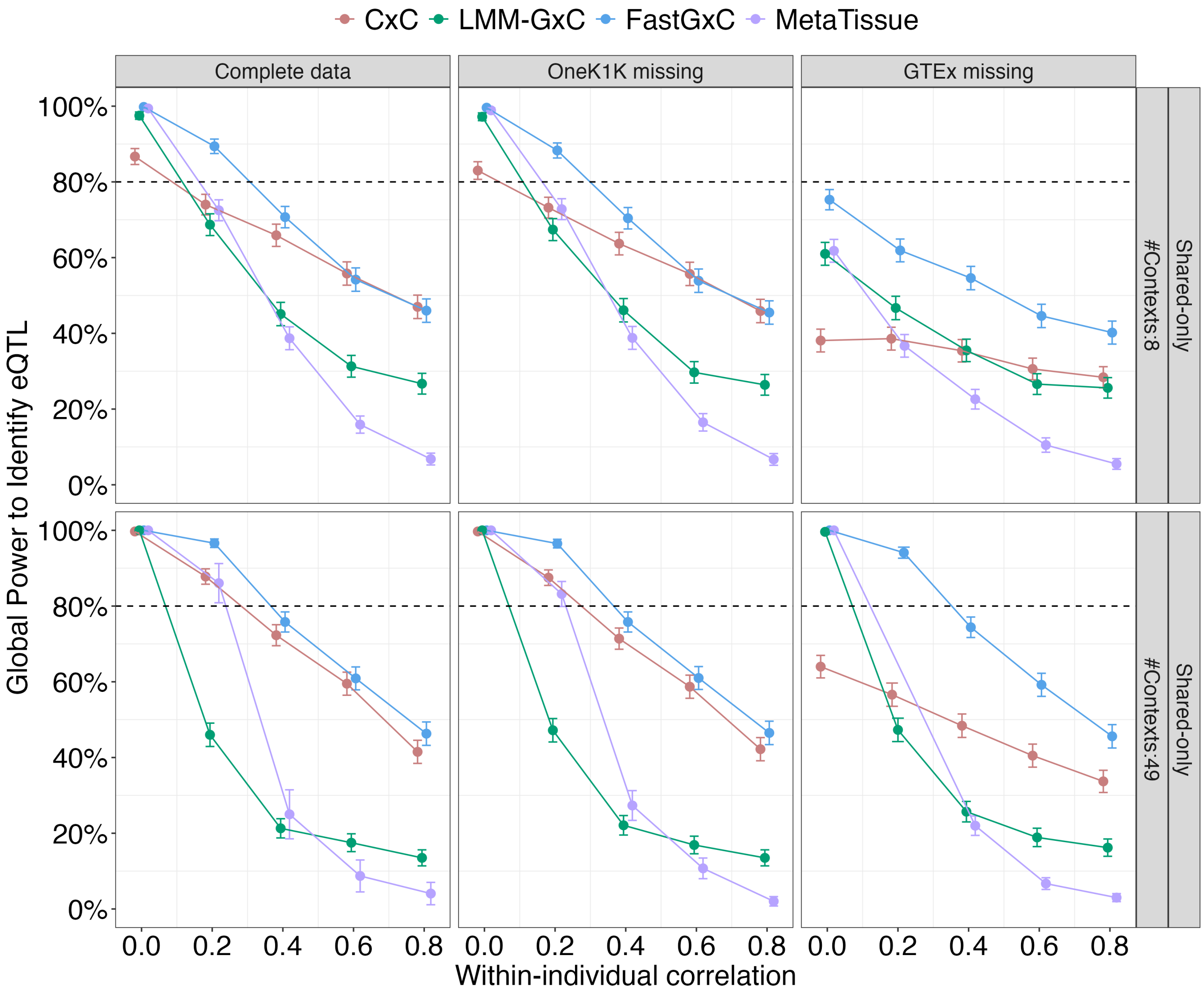


**Figure S5. Global power to identify an eQTL under no heterogeneity scenario.** Global power (y-axis) of each method (color) to identify a shared-only eQTL as a function of within-individual residual correlation of expression (x-axis), number of contexts (rows), and different missing data patterns(columns). Plot shows results for a sample size of 100 individuals. Results for sample size of 698 individuals are not plotted as all methods have nearly perfect power with current effect sizes. The LM-GxC method is not included in these plots because it is inflated under the null for testing this hypothesis.

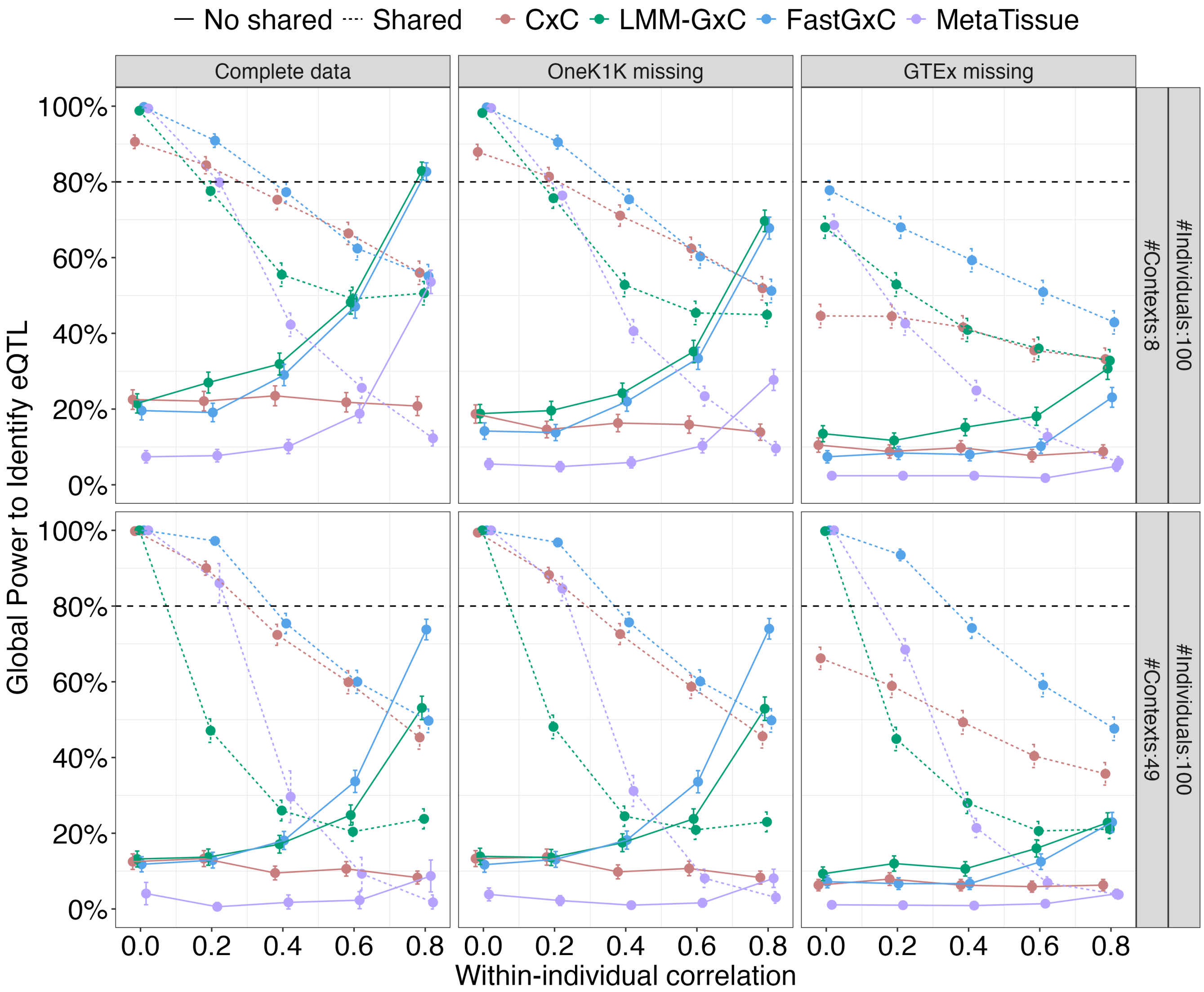


**Figure S6. Global power to identify an eQTL under a single-context heterogeneity scenario.** Global power (y-axis) of each method (color) to identify an eQTL under a single-context heterogeneity scenario as a function of within-individual residual correlation of expression (x-axis), presence of a context-shared effect (line type), number of contexts (rows), and different missing data patterns(columns). Plot shows results for a sample size of 100 individuals. Results for sample size of 698 individuals are not plotted as most methods have nearly perfect power with current effect sizes. Results for a two-context heterogeneity scenario look similar to these and are not plotted. The LM-GxC method is not included in these plots because it is inflated under the null for testing this hypothesis.

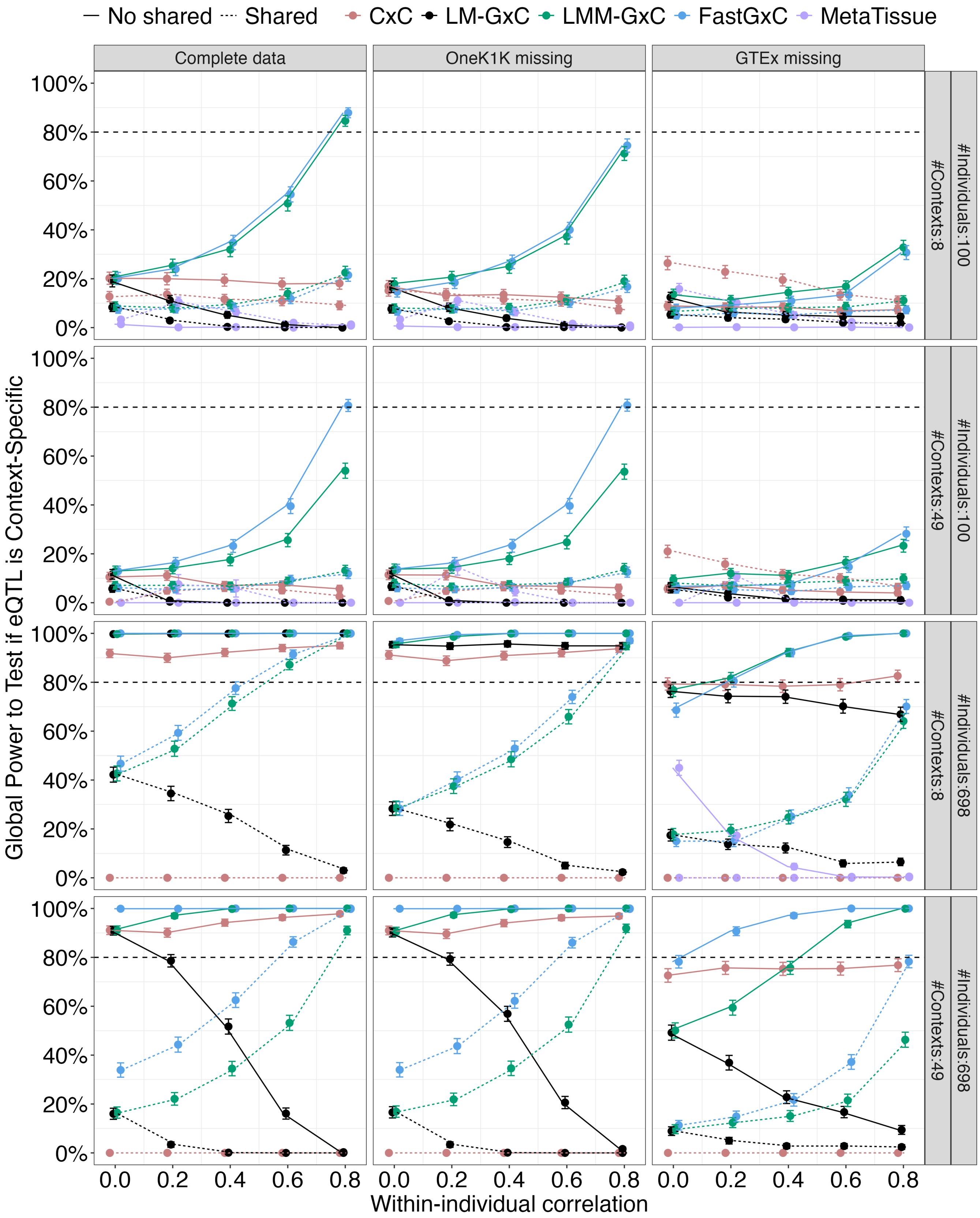


**Figure S7. Global power to test if eQTL is context-specific under a single-context heterogeneity scenario.** Global power (y-axis) of each method (color) to test if eQTL is context-specific under a single-context heterogeneity scenario as a function of within-individual residual correlation of expression (x-axis), presence of a context-shared effect (line type), number of contexts (rows), and different missing data patterns(columns).

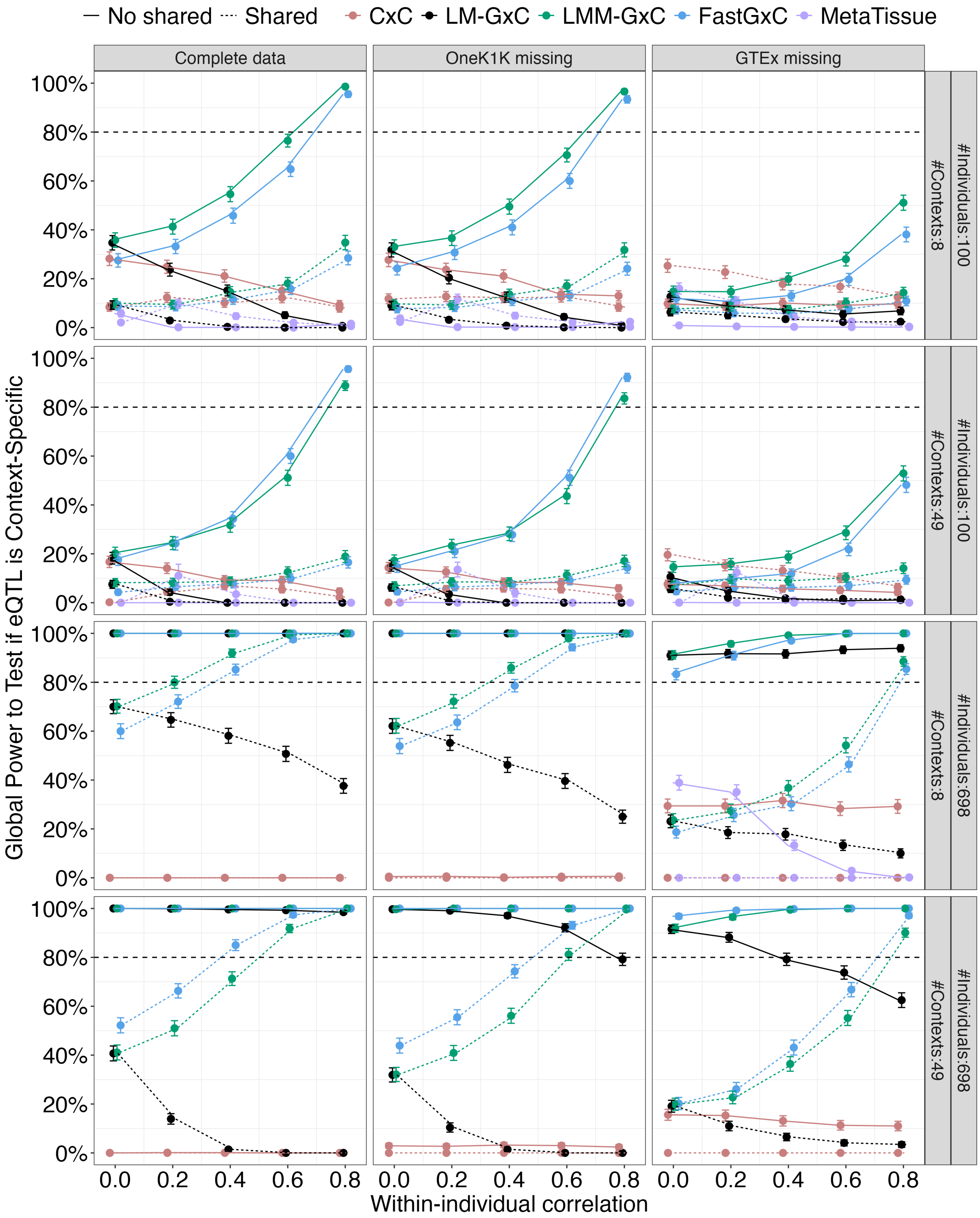


**Figure S8. Global power to test if eQTL is context-specific under a two-context heterogeneity scenario.** Global power (y-axis) of each method (color) to test if eQTL is context-specific under a two-context heterogeneity scenario as a function of within-individual residual correlation of expression (x-axis), presence of a context-shared effect (line type), number of contexts (rows), and different missing data patterns(columns).

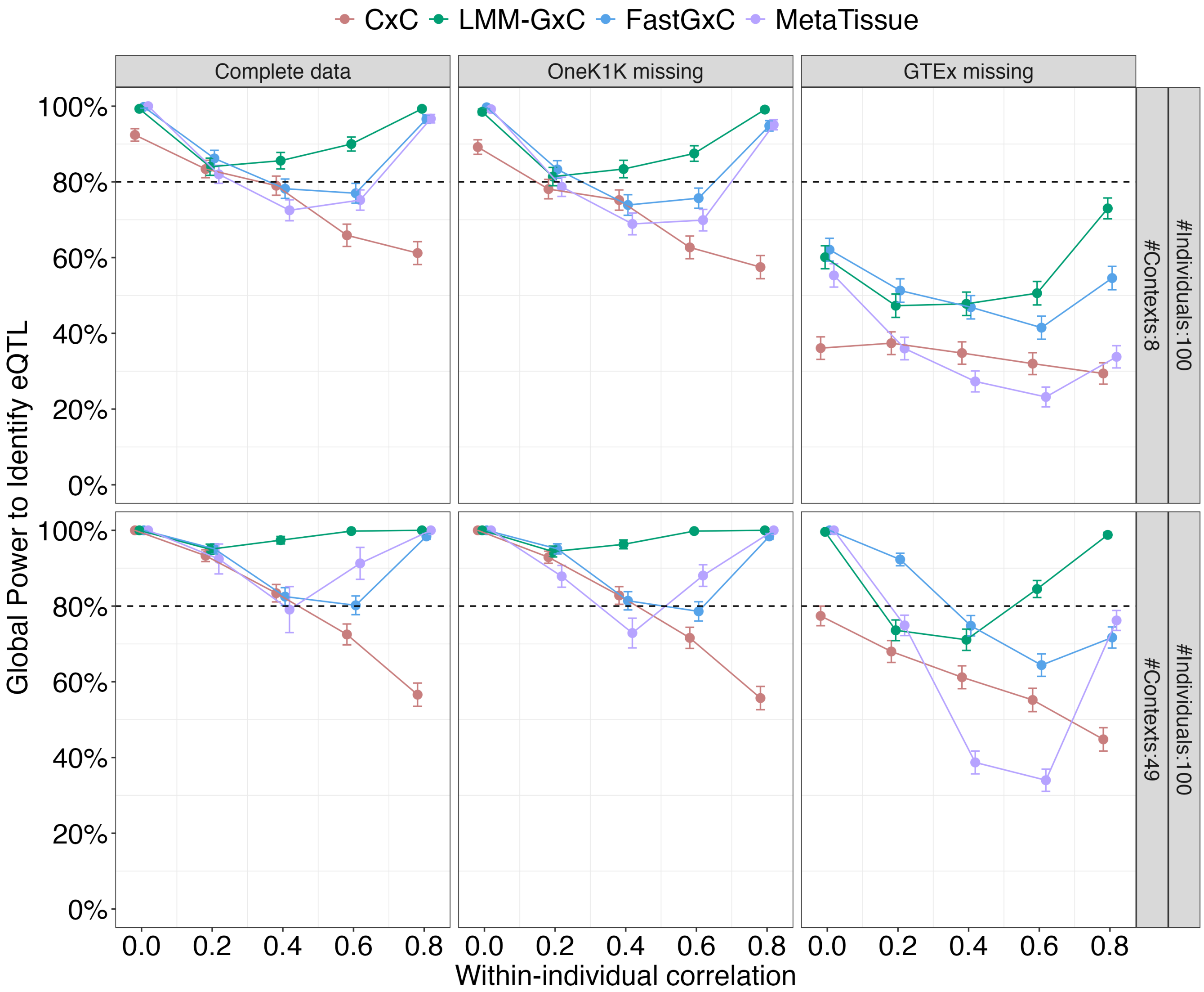


**Figure S9. Global power to identify an eQTL under an extensive heterogeneity scenario.** Global power (y-axis) of each method (color) to identify an eQTL under an extensive heterogeneity scenario as a function of within-individual residual correlation of expression (x-axis), number of contexts (rows), and different missing data patterns(columns). Plot shows results for a sample size of 100 individuals. Results for sample size of 698 individuals are not plotted as most methods have nearly perfect power with current effect sizes. The LM-GxC method is not included in these plots because it is inflated under the null for testing this hypothesis.

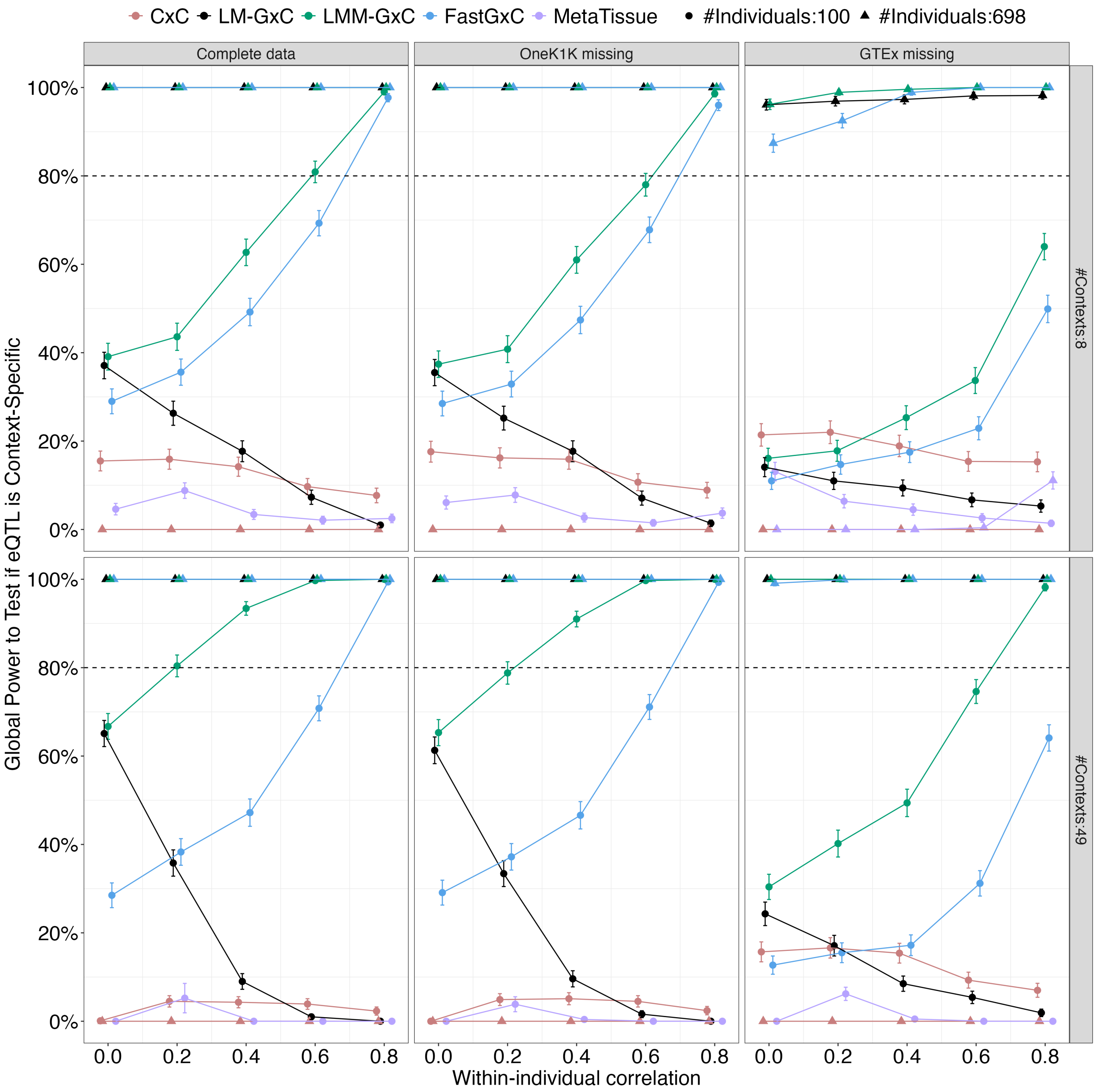


**Figure S10. Global power to test if eQTL is context-specific under an extensive heterogeneity scenario.** Global power (y-axis) of each method (color) to test if eQTL is context-specific under an extensive heterogeneity scenario as a function of within-individual residual correlation of expression (x-axis), number of contexts (rows) and individuals (shape), and different missing data patterns(columns).

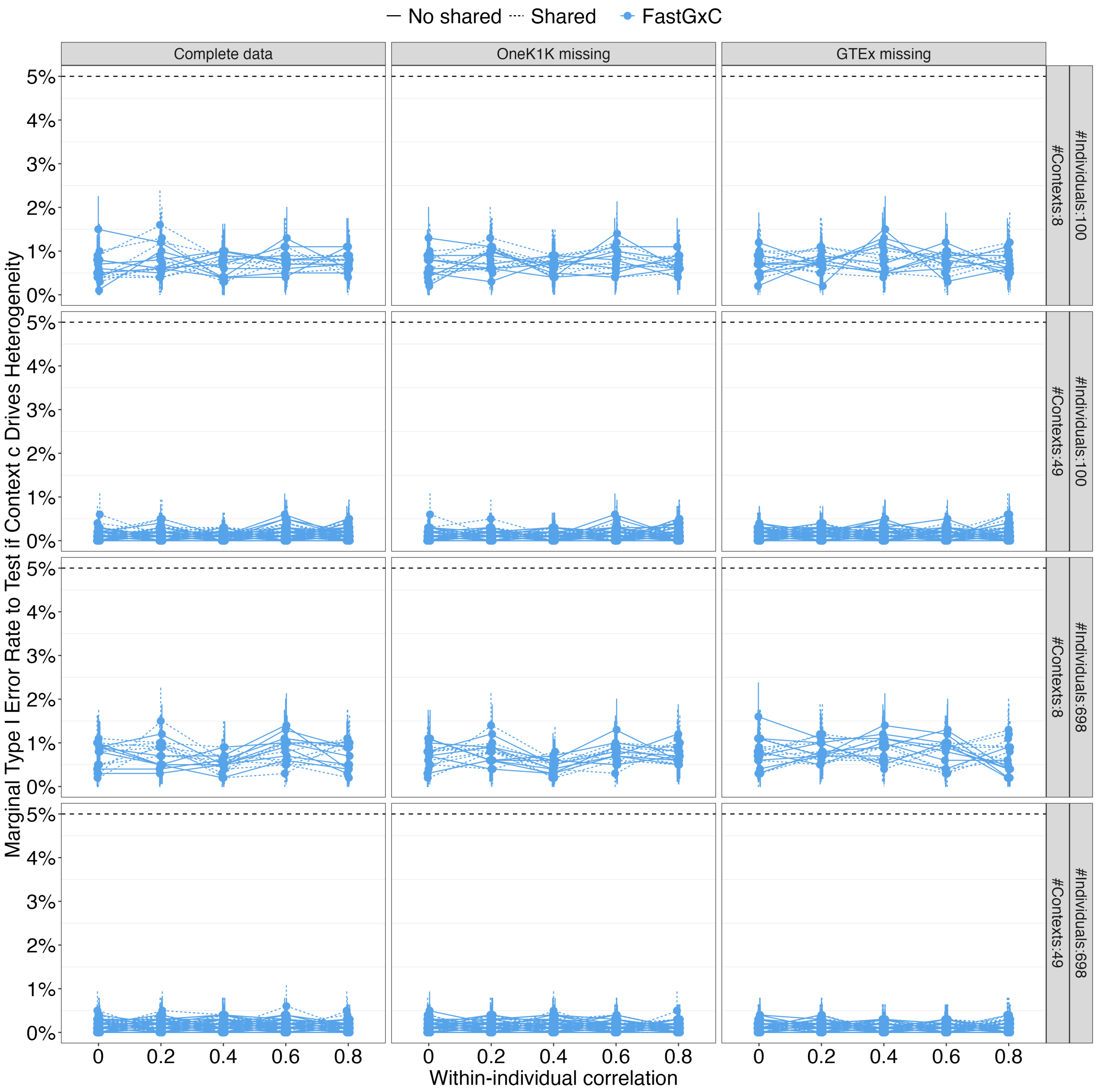


**Figure S11. FastGxC marginal type 1 error rate to test if a specific context c drives the heterogeneity.** Marginal type 1 error rate as a function of within-individual residual correlation of expression (x-axis), presence of a context-shared effect (line type), number of contexts and individuals (rows), and different missing data patterns(columns).

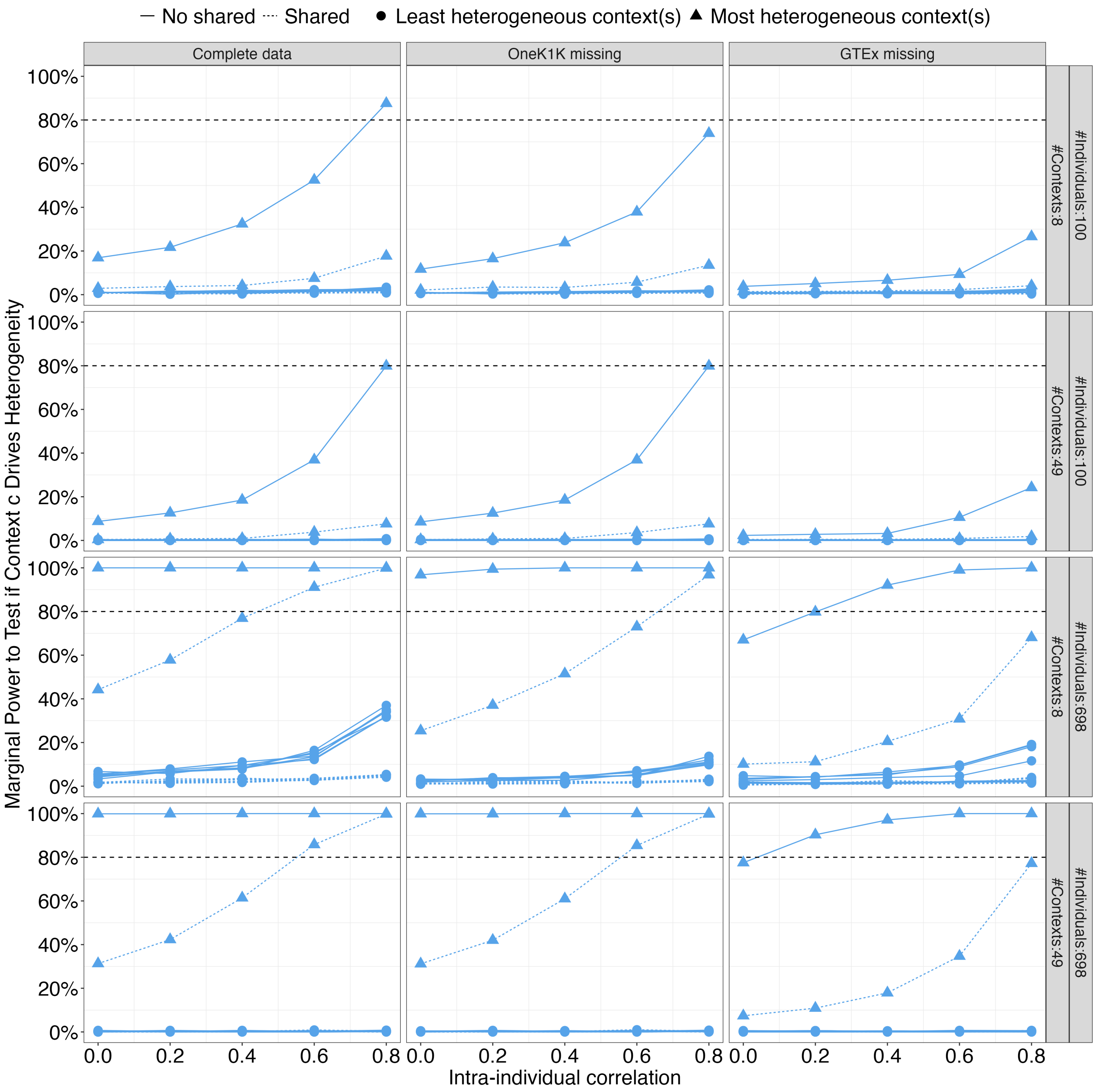


**Figure S12. FastGxC marginal power to test if a specific context c drives the heterogeneity under a single-context heterogeneity scenario.** FastGxC marginal power in the single-context heterogeneity case as a function of within-individual residual correlation of expression (x-axis), presence of a context-shared effect (line type), number of contexts and individuals (rows), and different missing data patterns(columns). Triangles represent contexts with a simulated (context-specific) eQTL and circles represent contexts without an (context-specific) eQTL (when a shared effect is present [linetype]).

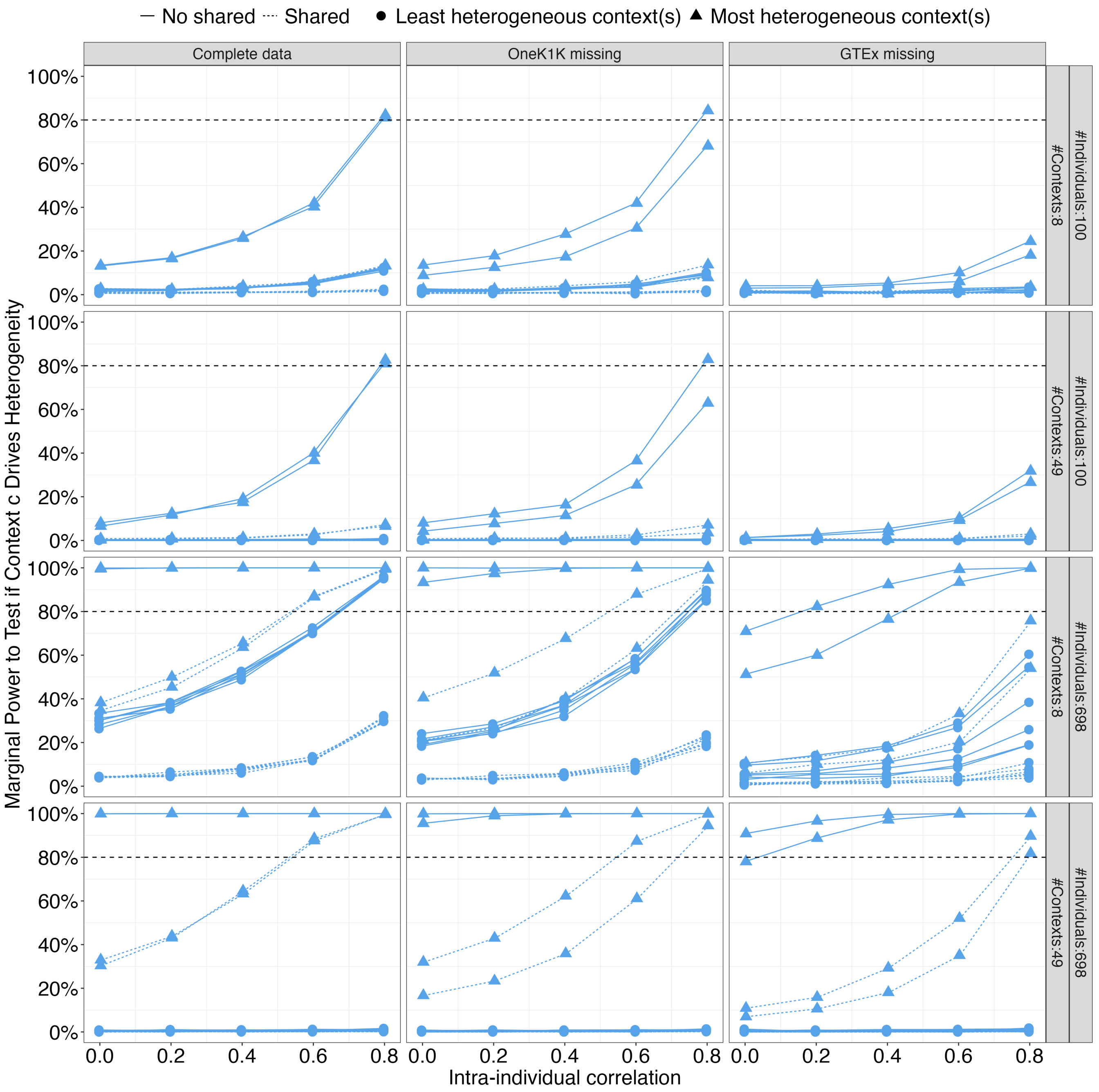


**Figure S13. FastGxC marginal power to test if a specific context c drives the heterogeneity under a two-context heterogeneity scenario.** FastGxC marginal power in the two-context heterogeneity case as a function of within-individual residual correlation of expression (x-axis), presence of a context-shared effect (line type), number of contexts and individuals (rows), and different missing data patterns(columns). Triangles represent contexts with a simulated (context-specific) eQTL and circles represent contexts without an (context-specific) eQTL (when a shared effect is present [linetype]).

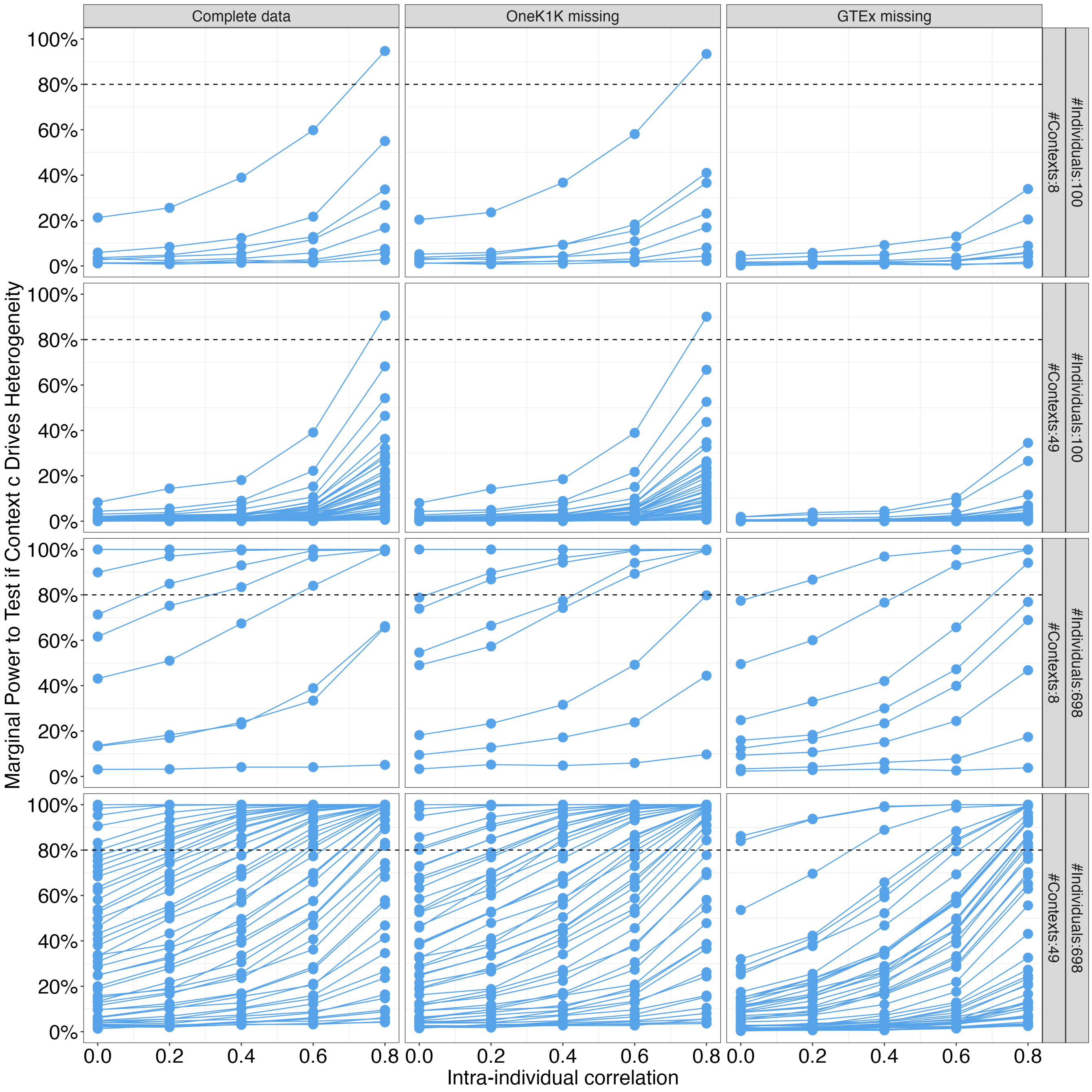


**Figure S14. FastGxC marginal power to test if a specific context c drives the heterogeneity under an extensive heterogeneity scenario.** FastGxC marginal power in the extensive heterogeneity case as a function of within-individual residual correlation of expression (x-axis), number of contexts and individuals (rows), and different missing data patterns(columns).

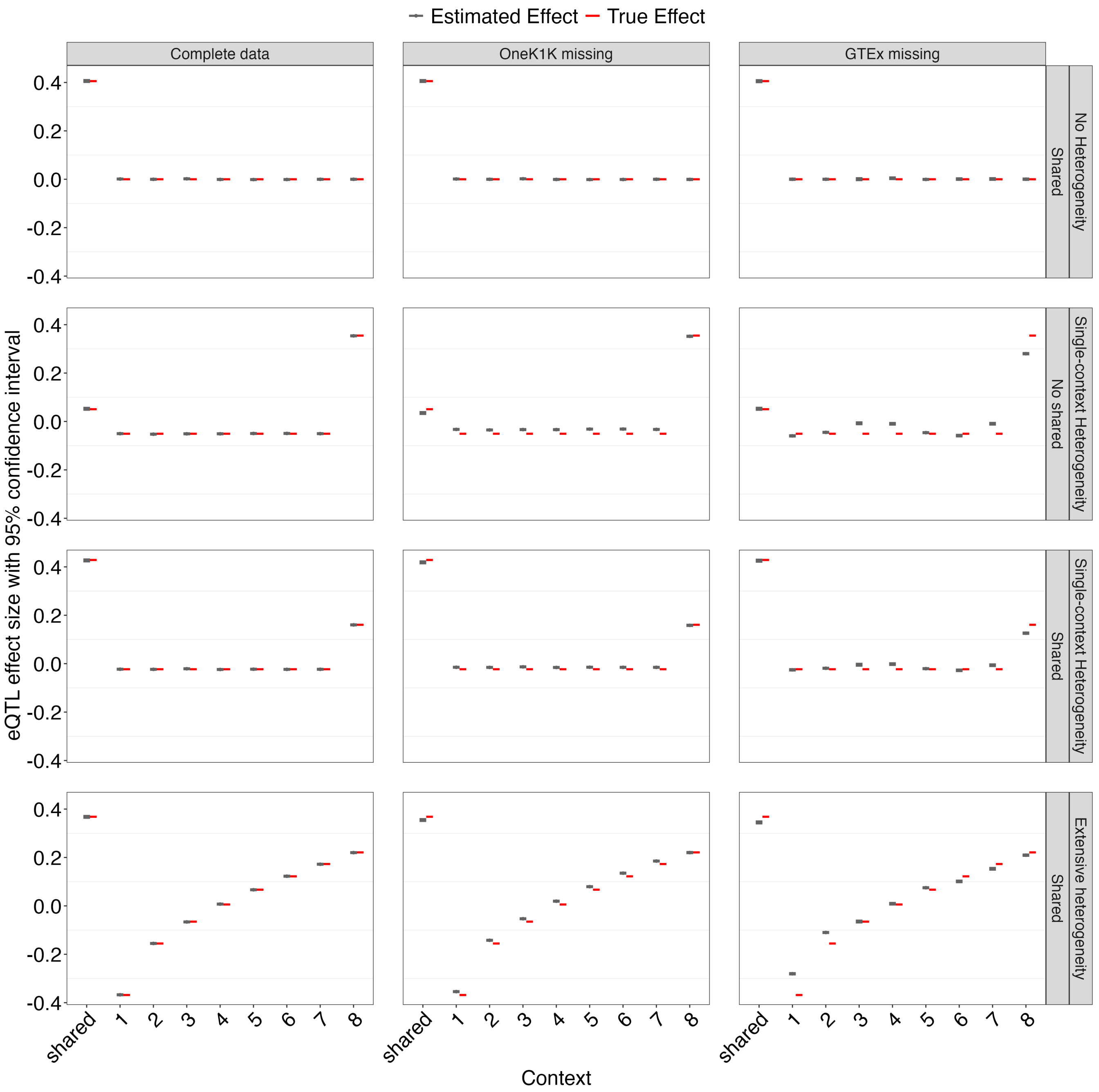


**Figure S15. Ability of FastGxC to estimate shared and specific eQTL effects across contexts with varying heterogeneity.** FastGxC effect size estimates with 95% confidence interval (grey) and true simulated effect size (red) in each of 8 contexts (x-axis). Effect sizes are shown across 4 different heterogeneity scenarios (rows), different missing data patterns (columns), and intra-individual correlation set to 0.8 for 698 individuals.

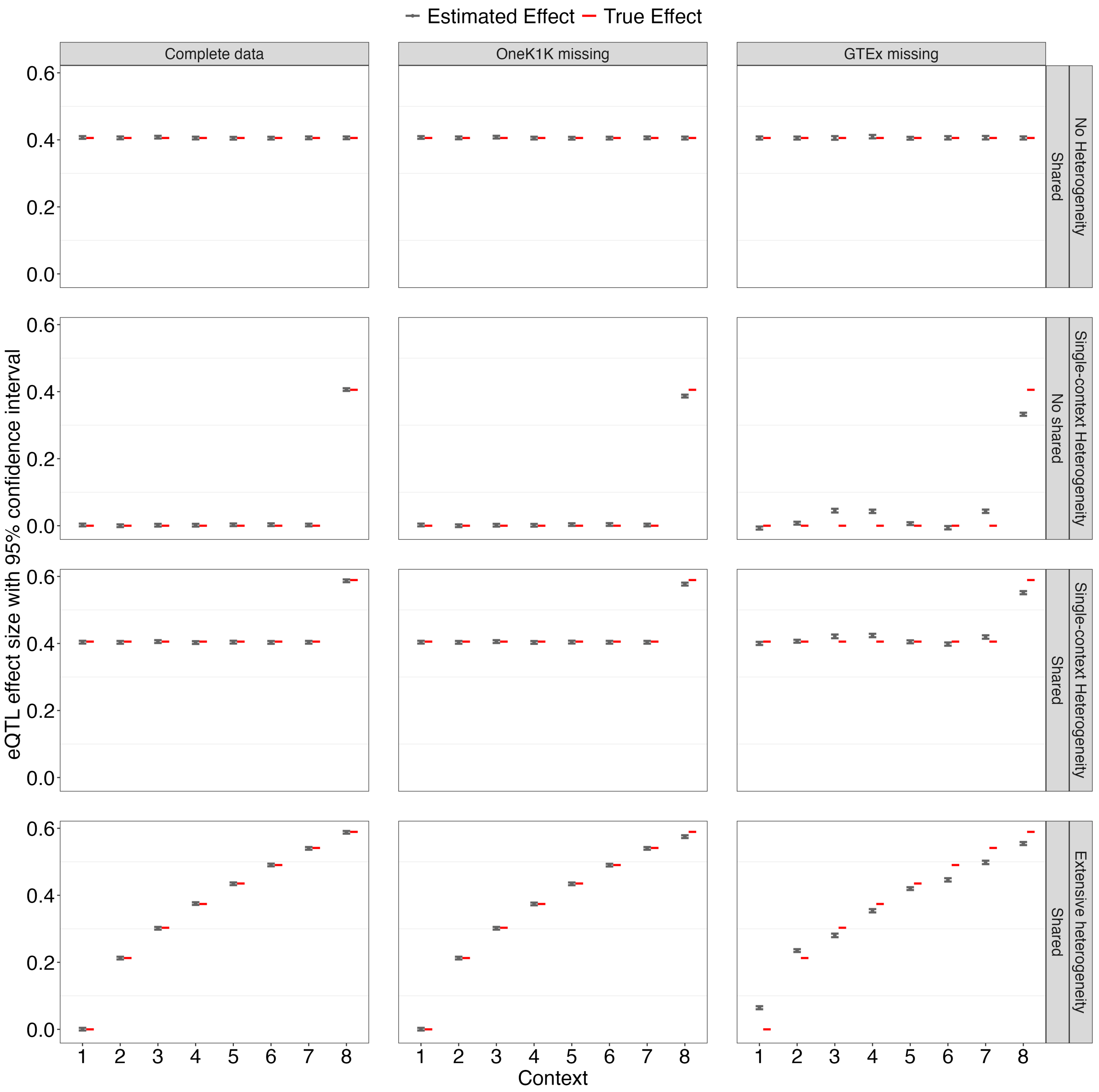


**Figure S16. Ability of FastGxC to estimate total eQTL effects across contexts with varying heterogeneity.** FastGxC effect size estimates with 95% confidence interval (grey) and true simulated effect size (red) in each of 8 contexts (x-axis). Effect sizes are shown across 4 different heterogeneity scenarios (rows), different missing data patterns (columns), and intra-individual correlation set to 0.8 for 698 individuals.

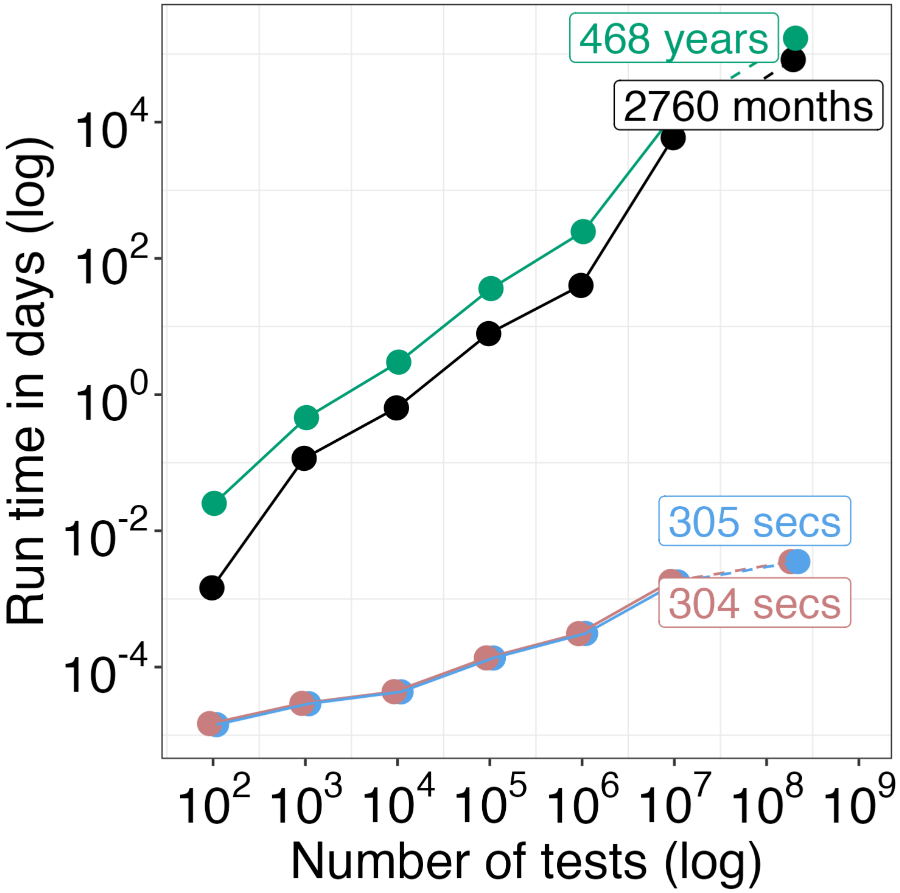


**Figure S17. Run time of each method in a simulated scenario with 1,000 individuals.**

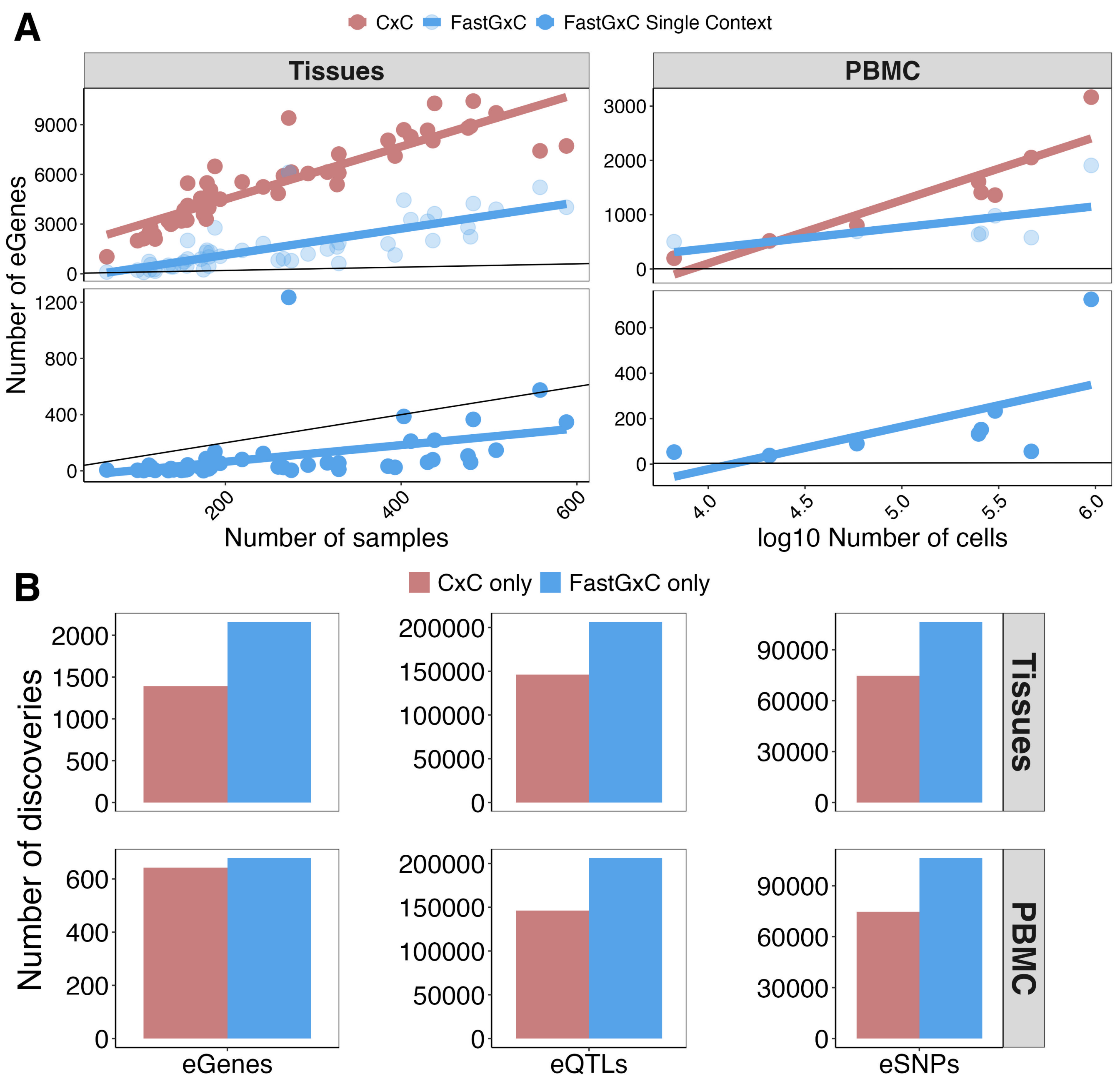


**Figure S18. Comparison of FastGxC and CxC discoveries in tissues and PBMC cell types. A.** Spearman correlation between number of samples (GTEx) or cell types (PBMC) and number of specific or single context-specific (opacity) eGenes per context and method (color). Solid black line shows y=x. **B.** Number of discoveries that are mapped uniquely by each method (color).

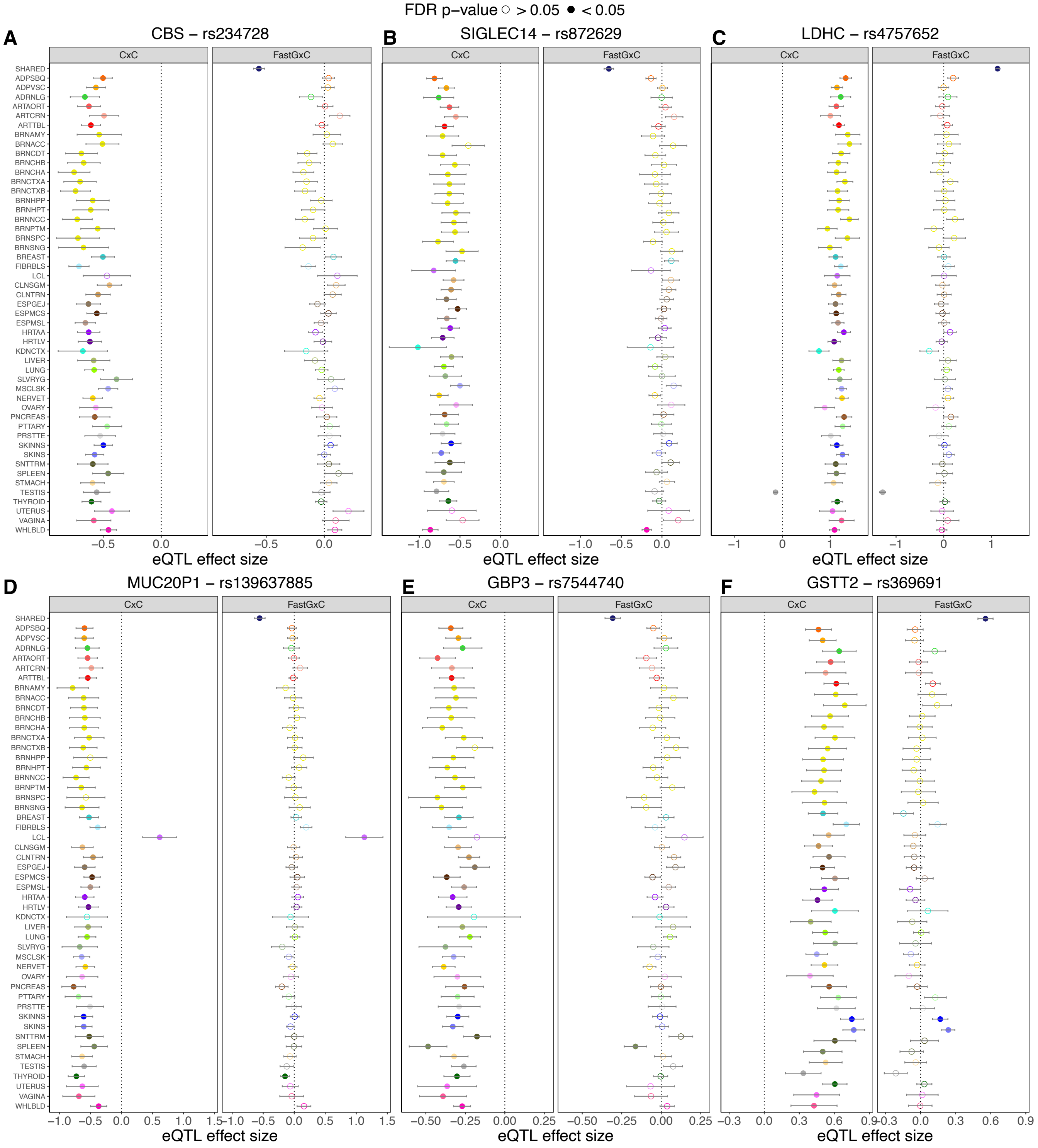


**Figure S19. Examples of eQTL mapped by FastGxC in GTEx.** Each panel represents a gene. Each dot shows the effect size estimates from CxC (L) and FastGxC (R) for a single tissue (color). Solid dots indicate a significant effect at 5% FDR.

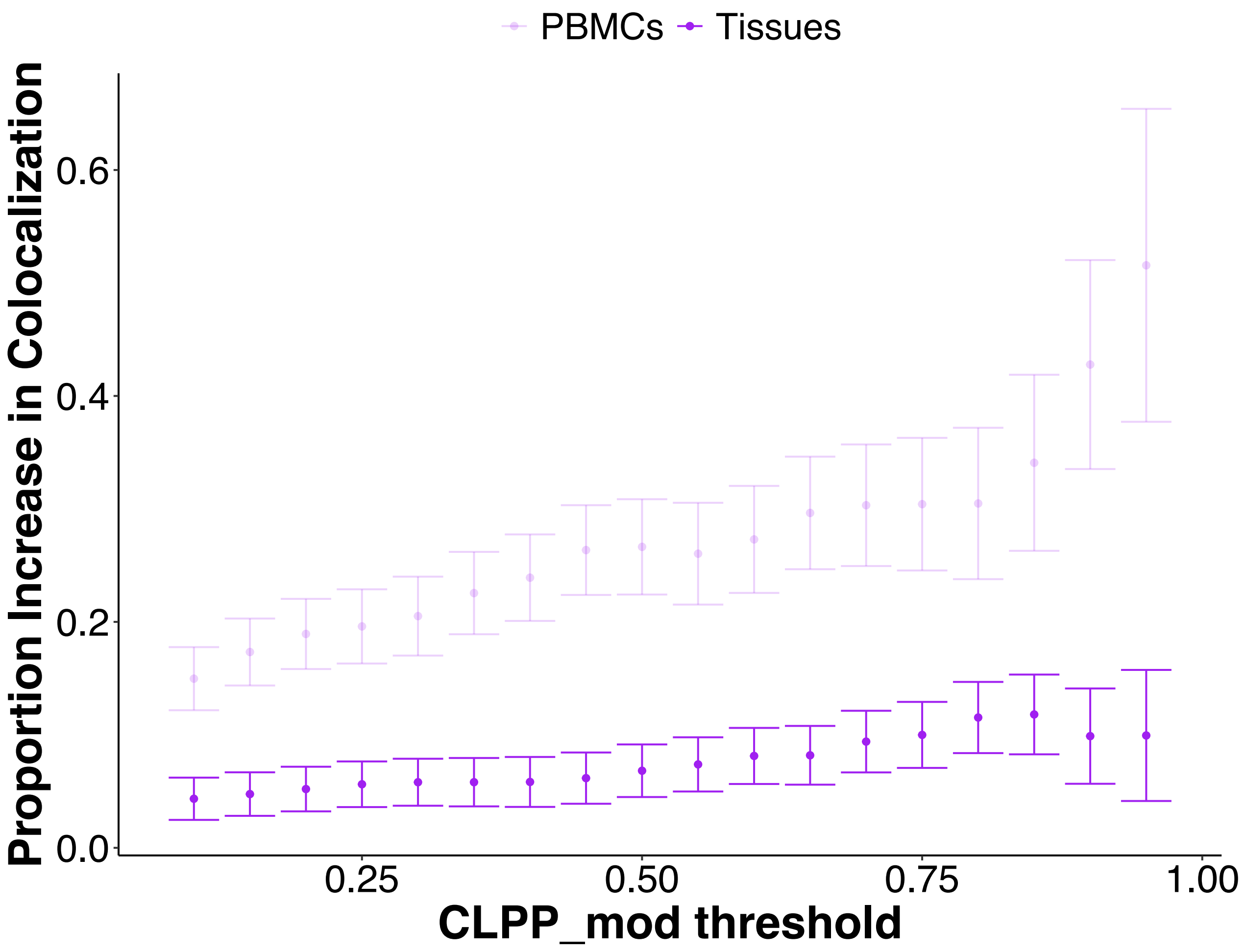


**Figure S20. Proportional increase of colocalizations FastGxC adds across varying $CLPP_{mod}$ thresholds.** Dark purple signifies proportional increase in tissues while lighter purpole is increase for PBMCs. Bars around points represent confidence intervals dependent on total number of colocalizations found at that $CLPP_{mod}$ threshold.

**Table S1. Comparison of existing methods developed to identify context-specific eQTLs.** Methods are grouped by those that jointly analyze data across contexts and context-by-context approaches that perform eQTL mapping followed by post hoc examination of summary statistics.

| Method | Outcome-Model | Handles Repeated Sampling? | Direct GxC test? | Predefined Contexts? | Linear GxC? |
|---|---|---|---|---|---|
| **Jointly analyze data across contexts** | | | | | |
| **Pseudo-bulk** | | | | | |
| LM-GxC [81] | TC-LM | No | Yes | Yes | Yes |
| LMM-GxC | TC-LM | Yes | Yes | Yes | Yes |
| FastGxC | TC-LM | Yes | Yes | Yes[1] | Yes |
| **Cell level** | | | | | |
| CellRegMap [25] | TC-LM | Yes | Yes | Yes | Yes |
| Airqtl [29] | TC-LM | Yes | Yes | Yes | Yes |
| scDALI [27] | ASC-GLM | Yes | Yes | Yes | Yes |
| Airpart [26] | ASC-GLM | Yes | Yes | Yes[1] | Yes |
| DAESC [28] | ASC-GLM | Yes | Yes | Yes | Yes |
| GASPACHO [30] | TC-GPLVM | Yes | Yes | No[2] | No |
| SURGE [24] | TC-LM | Yes | Yes | No[3] | Yes |
| **Context-by-context eQTL mapping followed by post hoc test** | | | | | |
| **Pseudo-bulk** | | | | | |
| CxC [10, 12, 15] | TC-LM | No | No [4] | Yes[1] | Yes |
| jaxQTL [33] | TC-GLM | No | No[5] | Yes[1] | Yes |
| **Cell level** | | | | | |
| SAIGE-QTL [31] | TC-GMM | Yes [6] | No [4] | Yes[1] | Yes |

(G)L(M)M: (generalized) linear (mixed) model; GxC: genotype-by-context interaction; TC: total counts; ASC: allele-specific counts; GPLVM: Gaussian process latent variable model.
[1]Discrete only. [2]Inferred via Gaussian process. [3]Inferred via matrix factorization. [4]Single context significant at FDR$\leq$ 5%. [5](Not) shared by sign and/or magnitude. [6]Within but not across contexts.

**Table S2. Parameter values for different simulated scenarios.** Each row corresponds to a different combination of parameters.

| Scenario | Heterogeneity Type | $\rho$ | $h_1^2$ | $h_2^2$ | $h_3^2$ | $h_4^2$ | $h_5^2$ | $h_6^2$ | $h_7^2$ | $h_8^2$ |
|---|---|---|---|---|---|---|---|---|---|---|
| 1-5 | No heterogeneity - No shared | 0.0-0.8 | 0.00 | 0.00 | 0.00 | 0.00 | 0.00 | 0.00 | 0.00 | 0.00 |
| 6-10 | No heterogeneity - Shared | 0.0-0.8 | 0.05 | 0.05 | 0.05 | 0.05 | 0.05 | 0.05 | 0.05 | 0.05 |
| 11-15 | Single-context heterogeneity - No shared | 0.0-0.8 | 0.00 | 0.00 | 0.00 | 0.00 | 0.00 | 0.00 | 0.00 | 0.05 |
| 16-20 | Single-context heterogeneity - Shared | 0.0-0.8 | 0.05 | 0.05 | 0.05 | 0.05 | 0.05 | 0.05 | 0.05 | 0.10 |
| 21-25 | Two-context heterogeneity - No shared | 0.0-0.8 | 0.00 | 0.00 | 0.00 | 0.00 | 0.00 | 0.00 | 0.05 | 0.05 |
| 26-30 | Two-context heterogeneity - Shared | 0.0-0.8 | 0.05 | 0.05 | 0.05 | 0.05 | 0.05 | 0.05 | 0.10 | 0.10 |
| 31-35 | Extensive Heterogeneity | 0.0-0.8 | 0.00 | 0.01 | 0.01 | 0.02 | 0.03 | 0.04 | 0.04 | 0.05 |

$\rho$: intra-individual residual correlation; $h_c^2$: proportion of expression variability explained by eQTL in context $c$

**Table S3.** Colors and abbreviations for GTEx tissues and PBMC cell types.

| tissue | color_hex | abbreviation |
|---|---|---|
| Adipose_Subcutaneous | #FF6600 | ADPSBQ |
| Adipose_Visceral_Omentum | #FFAA00 | ADPVSC |
| Adrenal_Gland | #33DD33 | ADRNLG |
| Artery_Aorta | #FF5555 | ARTAORT |
| Artery_Coronary | #FFAA99 | ARTCRN |
| Artery_Tibial | #FF0000 | ARTTBL |
| Brain_Amygdala | #EEEE00 | BRNAMY |
| Brain_Anterior_cingulate_cortex_BA24 | #EEEE00 | BRNACC |
| Brain_Caudate_basal_ganglia | #EEEE00 | BRNCDT |
| Brain_Cerebellar_Hemisphere | #EEEE00 | BRNCHB |
| Brain_Cerebellum | #EEEE00 | BRNCHA |
| Brain_Cortex | #EEEE00 | BRNCTXA |
| Brain_Frontal_Cortex_BA9 | #EEEE00 | BRNCTXB |
| Brain_Hippocampus | #EEEE00 | BRNHPP |
| Brain_Hypothalamus | #EEEE00 | BRNHPT |
| Brain_Nucleus_accumbens_basal_ganglia | #EEEE00 | BRNNCC |
| Brain_Putamen_basal_ganglia | #EEEE00 | BRNPTM |
| Brain_Spinal_cord_cervical_c-1 | #EEEE00 | BRNSPC |
| Brain_Substantia_nigra | #EEEE00 | BRNSNG |
| Breast_Mammary_Tissue | #33CCCC | BREAST |
| Cells_Cultured_fibroblasts | #AAEEFF | FIBRBLS |
| Cells_EBV-transformed_lymphocytes | #CC66FF | LCL |
| Colon_Sigmoid | #EEBB77 | CLNSGM |
| Colon_Transverse | #CC9955 | CLNTRN |
| Esophagus_Gastroesophageal_Junction | #8B7355 | ESPGEJ |
| Esophagus_Mucosa | #552200 | ESPMCS |
| Esophagus_Muscularis | #BB9988 | ESPMSL |
| Heart_Atrial_Appendage | #9900FF | HRTAA |
| Heart_Left_Ventricle | #660099 | HRTLV |
| Kidney_Cortex | #22FFDD | KDNCTX |
| Liver | #AABB66 | LIVER |
| Lung | #99FF00 | LUNG |
| Minor_Salivary_Gland | #99BB88 | SLVRYG |
| Muscle_Skeletal | #AAAAFF | MSCLSK |
| Nerve_Tibial | #FFD700 | NERVET |
| Ovary | #FFAAFF | OVARY |
| Pancreas | #995522 | PNCREAS |
| Pituitary | #AAFF99 | PTTARY |
| Prostate | #DDDDDD | PRSTTE |
| Skin_Not_Sun_Exposed_Suprapubic | #0000FF | SKINNS |
| Skin_Sun_Exposed_Lower_leg | #7777FF | SKINS |
| Small_Intestine_Terminal_Ileum | #555522 | SNTTRM |
| Spleen | #778855 | SPLEEN |
| Stomach | #FFDD99 | STMACH |
| Testis | #AAAAAA | TESTIS |
| Thyroid | #006600 | THYROID |
| Uterus | #FF66FF | UTERUS |
| Vagina | #FF5599 | VAGINA |
| Whole_Blood | #FF00BB | WHLBLD |
| Shared | midnightblue | SHARED |

| cell_type | color_hex | abbreviation |
|---|---|---|
| Shared | midnightblue | SHARED |
| B | #660099 | B |
| CD4 | #AAEEFF | CD4 |
| CD8 | #0000FF | CD8 |
| cMono | #EEEE00 | CMONO |
| ncMono | #FFAA00 | NCMONO |
| pDC | #006600 | PDC |
| cDC | #99BB88 | CDC |
| NK | #FFAAFF | NK |

**Table S4. FastGxC mapped shared and specific eGenes in tissues and PBMCs.** FastGxC shared and specific eGenes from GTEx tissues and OneK1K+CLUES meta analyzed cell types are provided as a separate excel file, one sheet per study, with the following columns: eGene type (sh-eGene or sp-eGene), tissue or cell type, gene identifier.

**Table S5. EQTL enrichment in GWAS loci results.** Results from enrichment of GTEx and single-cell eQTLs from CxC and FastGxC in GWAS catalog loci are provided as a separate excel file with two sheets. The first sheet shows manual annotation of most likely relevant tissue(s) for GWAS catalog traits with the following columns: GWAS Trait, Most Likely Relevant Tissue(s). The second sheet shows enrichment results with the following columns: GWAS trait, method, tissue (GTEx) or cell type (PBMCs), enrichment odds ratio (OR), OR lower confidence interval, OR upper confidence interval, enrichment p-value.

**Table S6. Colocalization of FastGxC or CxC eQTLs with GWAS variants.** Colocalization results from tissue and cell type FastGxC and CxC eQTLs with GWAS hits are provided as a separate excel file with two sheets, one for GTEx and one for OneK1K+CLUES meta analysis, with the following columns: lead_snp, gwas_trait, tissue (GTEx) or celltype (PBMCs), study, method, gene, n_snps, clpp, clpp_mod.
